# Harvesting-induced evolution of collective risk-taking behavior and changes to the circadian system in a fish

**DOI:** 10.1101/622043

**Authors:** Valerio Sbragaglia, Jose Fernando López-Olmeda, Elena Frigato, Cristiano Bertolucci, Robert Arlinghaus

## Abstract

Intensive and trait-selective harvesting of fish and wildlife can cause evolutionary changes in a range of life-history and behavioural traits. These changes might in turn alter the circadian system both at behavioral and molecular levels, with knock-on effects on daily physiological processes and behavioural outputs. We examined the evolutionary impact of size-selective mortality on collective risk-taking behavior and the circadian system in a model fish species. We exposed zebrafish (*Danio rerio*) to either large or small size-selective mortality relative to a control over five generations, followed by eight generations during which harvesting halted to remove maternal effects. Large size-selective mortality typical of many fisheries and specialized predators targeting larger size classes decreased collective risk-taking behavior compared to controls. The opposite occurred in response to small size-selective mortality typical of specialized fisheries and most gape-limited predators. The evolutionary changes in risk-taking behavior were linked to daily activity rhythms in response to small size-selective mortality, while no changes were observed in response to large size-selective mortality. We also found changes in the molecular circadian core clockwork in response to both size selective mortality treatments. These changes disappeared in the clock output pathway, resulting in similar transcription profiles of both size-selected lines. The results suggest a switch downstream to the molecular circadian core clockwork, leading to overall similar daily activity patterns across selection lines. Our experimental harvest left an evolutionary legacy in collective risk-taking behavior and relatedly in the circadian system, both at behavioral and molecular levels. Changes to risk-sensitive behavior of exploited organisms can have far-reaching consequences for how space and time is used and may also affect catchability and natural predation.

## INTRODUCTION

Harvesting is among the five major forms of human-induced rapid environmental change (Díaz *et al.* 2020). Harvesting differs from many other forms of natural predation by primarily targeting adult individuals – a size class that typically experiences limited natural mortality (Darimont *et al.* 2015). Intensive and trait-selective harvesting thus can shift the fitness landscape and foster evolutionary adaptations at a rate and speed that is rarely experienced in the evolutionary history of many animal populations (Palumbi 2001; Jørgensen *et al.* 2007; Allendorf & Hard 2009; Sih, Ferrari & Harris 2011).

Empirical and modeling studies on the topic of fisheries-induced evolution have suggested intensive fishing fosters evolution of a fast life history that is characterized by elevated reproductive investment, reduced age and size at maturation and reduced post maturation growth and longevity (Jørgensen *et al.* 2007; Heino, Díaz Pauli & Dieckmann 2015). Fisheries-induced evolution might also affect behavioral and physiological traits, but they are much less studied compared to life-history adaptations (Uusi-Heikkilä *et al.* 2008; Heino, Díaz Pauli & Dieckmann 2015; Hollins *et al.* 2018).

The proximate mechanisms governing fisheries-induced evolution of behavior, as well as other traits, are difficult to disentangle in the wild because phenotypic changes observed in time series (e.g., changes in average age at maturation) can be masked by phenotypic plasticity, i.e., non-genetic changes in trait expression (Heino, Díaz Pauli & Dieckmann 2015). One means to single out the cause-and-effect of fishing is conducting selection experiments with model species in the laboratory (Conover & Baumann 2009; Díaz Pauli & Heino 2014). Several laboratories have investigated the evolutionary effects of size-selective harvesting on behavior by means of selection experiments with different fish species (e.g. Walsh *et al.* 2006; Uusi-Heikkilä *et al.* 2015; Díaz Pauli *et al.* 2019). This line of research has largely focused on evolutionary changes of life-history traits and additionally has started to study harvesting-induced changes in individual personality traits, such as boldness. Changes in individual behavioural traits may have strong repercussions for affecting collective behavioural traits (Jolles, King & Killen 2020). A focus on collective behaviour is crucial to fully understand the evolutionary changes in response to fishing or other forms of natural mortality because shoaling is a key driver of the capture process (e.g. Tenningen *et al.* 2016; Thambithurai *et al.* 2018) and has also strong adaptive value to reduce exposure to natural predators (Pitcher 1986; Sumpter 2010). More work is needed to understand whether and to what degree intensive size-selective mortality can affect individual and collective behavioural traits in fish and other species (Arlinghaus *et al.* 2017).

Size-selective harvesting can affect behavioral traits through at least two mechanisms (Fig. 1). First, following the pace-of-life syndrome (Réale *et al.* 2010), behavioural and life-history traits are often correlated along a fast-to-slow continuum. Fast life-history traits – those trait combinations favored under intensive and large size-selective harvesting pressure (e.g. Jørgensen & Holt 2013; Andersen, Marty & Arlinghaus 2018) - are expected to covary and co-evolve with an increase of risk-taking behavior (Wolf *et al.* 2007; Réale *et al.* 2010; Fig. 1). The argument is that acquiring the resources needed to maintain a fast life-history (i.e. rapid juvenile growth or high reproductive investment) favors risky behaviors such as feeding in the presence of predators. Under such scenario, harvesting-induced evolution would predict an increase of risk-taking behavior. Yet, recent refinements of the pace-of-life syndrome hypothesis suggest that specific ecological contexts (e.g. variation in natural mortality risk) could systematically break apart the genetic correlation of life-history and behavioural traits (Dammhahn *et al.* 2018; Montiglio *et al.* 2018). Indeed, observational (Polverino *et al.* 2018) and theoretical (Andersen, Marty & Arlinghaus 2018; Claireaux, Jorgensen & Enberg 2018) studies in fish have suggested that the covariance of life-history and risk-taking behavioural traits may no longer exist in response to mortality-based selection pressures. It is thus equally plausible that intensive and size-selective harvest may foster the evolution of both a fast life history and a decrease of risk-taking behavior (Andersen, Marty & Arlinghaus 2018).

**Figure 1.**
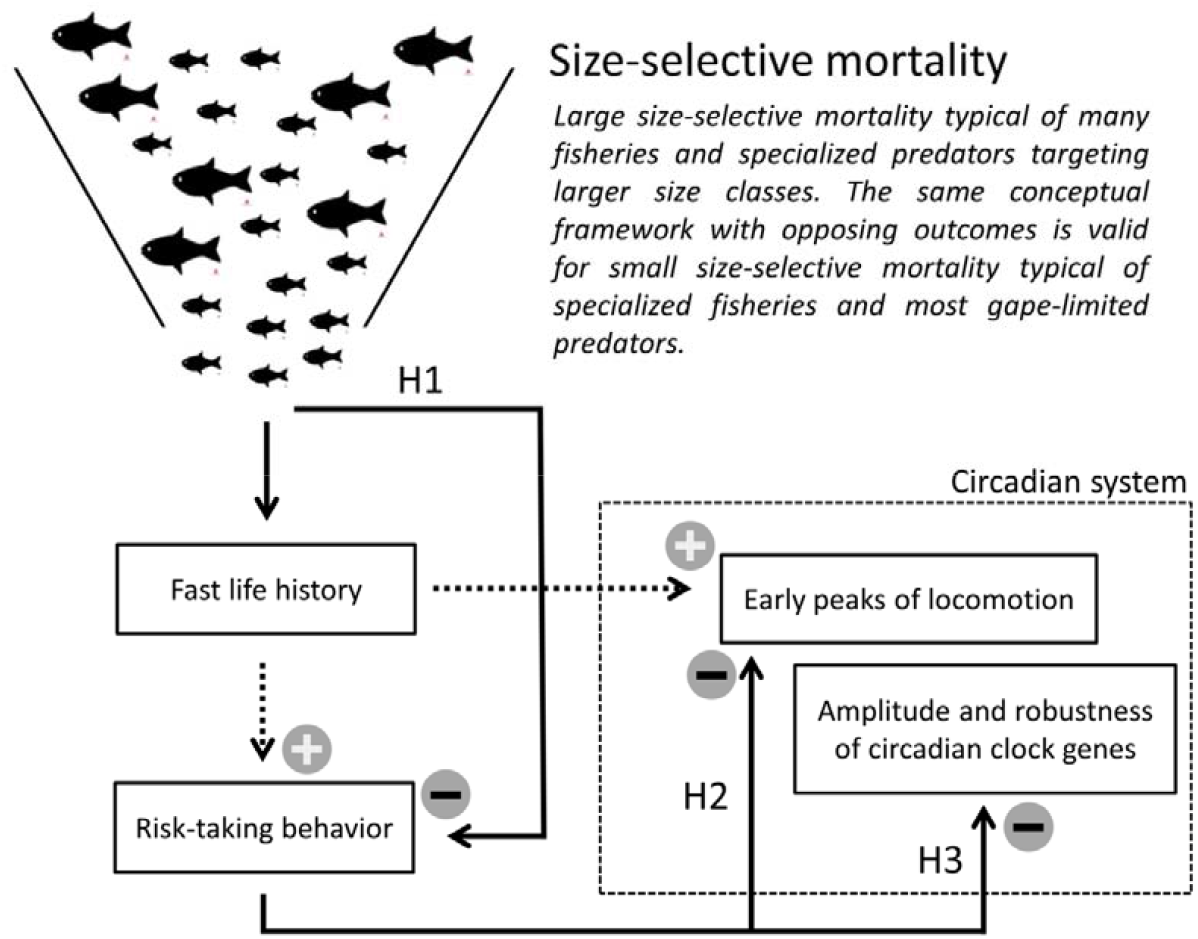
The conceptual framework of the putative mechanisms driving evolution of risk-taking behavior and circadian system in response to size-selective harvesting. Large size-selective harvesting triggers the evolution of a fast life history, which in turn can coevolve with an increase of both risk-taking behavior and early peaks of locomotor activity. However, a plausible scenario is that harvesting disrupts the correlation of life-history with other behavioral traits (dotted arrows). A further possibility is that large-size selective harvesting triggers the evolution of a decrease risk-taking behavior through a correlated selection response with size-at-age (H1). Consequently, a decrease of risk-taking behavior may affect the circadian system by triggering a decrease of early locomotor activity rhythms (H2), and reducing amplitude and robustness of circadian clock gene expression (H3). The same framework with opposite prediction is valid for small size-selective harvesting. For more details, see the main text.

Second, fish behavioral traits can evolve in response to correlated selection responses linked to size-selective mortality due to a systematic positive correlation of specific behaviors, growth rate and size-at-age (Biro & Post 2008; Biro & Sampson 2015; Klefoth *et al.* 2017). Specifically, behavioral traits related to resource acquisition, such as risk-taking behavior during foraging (boldness; Reale *et al.* 2007), increase food intake and thereby elevate growth rate and affect size-at-age (Enberg *et al.* 2012). In scenarios where the larger individuals have a higher risk of mortality due to fishing or perhaps specialized natural predators, selective removal of these larger, faster growing individuals might also indirectly favor a decrease of risk-taking behavior (Biro & Post 2008). Consequently, selective harvesting of large fish might evolutionarily favor a decrease of risk-taking behavior (Andersen, Marty & Arlinghaus 2018; Claireaux, Jorgensen & Enberg 2018; Fig. 1).

Evolutionary changes in behavioral traits can in turn affect the circadian system (Fig. 1). Recent studies have reported a relationship between copying style (or animal personality), particularly risk-taking behavior and the circadian system in zebrafish, *Danio rerio* (Rey *et al.* 2013; Tudorache *et al.* 2018). The circadian system is widely conserved phylogenetically (Panda, Hogenesch & Kay 2002) and controls daily rhythms of physiology and behavior (Dunlap, Loros & DeCoursey 2004). Thereby, circadian rhythms fulfill an adaptive function helping the organism to anticipate and maintain synchronization to external 24-h environmental cycles (Kronfeld-Schor & Dayan 2003; Idda *et al.* 2012; Cowan, Azpeleta & López-Olmeda 2017). Functional circadian clocks are essential for precise timing of activity, thereby helping the organism to avoid or minimize risk of predation while maximizing foraging (DeCoursey, Walker & Smith 2000).

Evolutionary changes of the circadian system in exploited populations may also occur due to the interplay between the evolutionary responses to life history and behavior (Fig. 1). Experimental evolution in *Drosophila melanogaster* indicated that selection for slower-running (faster-running) circadian clocks could result in slower (faster) rates of developmental processes, which in turn can delay (advance) development time (Nikhil & Sharma 2017 and references therein). Moreover, the clock gene *clock* (i.e., a gene that is part of the core transcription-translation feedback loop of the molecular circadian clockwork in fish; Vatine *et al.* 2011) is mainly mapped in regions related to life history traits in salmonids (Leder, Danzmann & Ferguson 2006; Paibomesai *et al.* 2010). This suggests that life history changes induced by harvesting could coevolve with the circadian clock (e.g., Leder, Danzmann & Ferguson 2006; Nikhil & Sharma 2017). Relatedly, proactive (i.e. more risk taking) zebrafish have shown more robust diurnal rhythms at molecular, endocrine and behavioural level than reactive ones (Tudorache *et al.* 2018). Therefore, harvesting induced evolution of risk-taking behavior can also be expected to affect the circadian system both at behavioral and molecular levels (Fig. 1). Studying both levels is relevant because phenotypic changes might not be mirrored at the genetic level, as recently demonstrated in a harvesting-induced evolution experiment using Atlantic silversides, *Menida menida* (Therkildsen *et al.* 2019).

We take advantage of a long-term harvesting experimental system based on zebrafish (Uusi-Heikkilä *et al.* 2015). The zebrafish selection lines were exposed to strong directional harvest selection (a 75% per-generation harvest rate) acting on either large (large-harvested line; a common scenario in many fisheries worldwide and in presence of predators where large individuals are selectively captured) or small (small-harvested line; a common scenarios in specific fisheries or in the presence of gape-limited predators that preferentially feed on the smaller size classes) fish, relative to a control line harvested randomly with respect to body size (Uusi-Heikkilä *et al.* 2015). Previous research has shown that the large-harvested line evolved a faster life history characterized by smaller adult length and weight, higher relative fecundity and elevated senescence compared to controls (Uusi-Heikkilä *et al.* 2015; see Fig. S1). By contrast, the small-harvested line showed a slower life history compared to controls, characterized by reduced reproductive investment and no change in adult length compared to the control line (Uusi-Heikkilä *et al.* 2015). Other previously documented changes include strong evidence for genetic and genomic changes, i.e., the differences among the selection lines were indicative of evolutionary and not just phenotypic adaptations (Uusi-Heikkilä *et al.* 2015; Uusi-Heikkilä *et al.* 2017). Size-selection occurred during the first five generations, but the size-selected lines maintained their key life-history adaptations after harvesting halted for up to eight generations (Uusi-Heikkilä *et al.* 2015; Sbragaglia *et al.* 2019b), indicating that harvesting-induced evolution of life-history traits did not recover to pre-harvesting levels (Fig. S1). The fixation of the harvesting-induced changes in life history after harvesting halted is a precondition to allow a comparison among the selection lines in terms of harvesting-induced evolution of behavior and the circadian system. Addressing this question was the aim of the present research (Fig. 1).

We predicted (H1 in Fig. 1) that evolution of collective risk-taking behavior is governed by a resource acquisition mechanism in a way that faster growth rate is correlated with an increase of risk-taking behaviour (Enberg *et al.* 2012). Therefore, the selective removal of the larger, faster-growing individuals is predicted to result in the evolution of decreased collective risk-taking behavior in the large-harvested line (i.e., elevated shyness or timidity; Andersen, Marty & Arlinghaus 2018), and the opposite was expected for the small-harvested line. Second, we predicted that a decrease of collective risk-taking behavior in the large-harvested line triggers changes in the circadian system both at behavioral and molecular levels (Rey *et al.* 2013; Tudorache *et al.* 2018). If this is true, we expected the absence of early peaks of locomotor activity (H2 in Fig.1), because a decrease of risk-taking behavior has been associated with the absence of clear peaks of activity in zebrafish (Tudorache *et al.* 2018). Again, the opposite response was expected for the small-harvested line. Third, at the molecular level, we expected that the large-harvested line lost circadian rhythmicity of clock genes (H3 in Fig.1), because a decrease of risk-taking behavior has been associated with a loss of robustness and amplitude in clock genes expression (here quantified as circadian rhythmicity; Tudorache *et al.* 2018). Following well-established knowledge of the functioning of the zebrafish circadian system (e.g. Vatine *et al.* 2011; Idda *et al.* 2012), we focused our investigation on carefully selected genes related to fundamental properties of the circadian system functioning, such as the core transcriptional-translational feedback loop (Dunlap, Loros & DeCoursey 2004; Vatine *et al.* 2011), light-inducible genes (genes directly activated by light that can interfere with the functioning of the core feedback loop; Tamai, Young & Whitmore 2007; Vatine *et al.* 2011), circadian clock output (genes related to mechanisms downstream the core feedback loop), and clock controlled genes related to energy balance.

## MATERIAL AND METHODS

### Selection lines

Our experimental system consisted of wild-collected zebrafish from West Bengal in India, sampled with a range of fishing gears (seine, cast nets and dip nets). The parental wild-collected population was experimentally harvested as explained in detail elsewhere (Uusi-Heikkilä *et al.* 2015). Each selection line was replicated twice for a total of six selection lines, similar to a landmark study about the outcomes of experimental size-selective harvesting in Atlantic silversides (Conover & Munch 2002; Therkildsen *et al.* 2019). Zebrafish were exposed to size selection only during the first five generations and then harvesting halted for several further generations (Uusi-Heikkilä *et al.* 2015) to remove maternal effects and study evolutionary outcomes in a common-garden setting.

Zebrafish selection lines for the present experiment where from F_13_, eight generations after harvesting halted where the lines still showed and maintained key life-history adaptations where the large-harvested line showed a fast life-history (e.g., elevated reproductive investment and reduced post maturation growth) and the small-harvested line signs of a slow fast-history (Fig. S1; Uusi-Heikkilä *et al.* 2015). Fish were reared in groups under *ad libitum* feeding and maintained under the following conditions: water temperature at 26 ± 0.5 °C; photoperiod at 12:12 h LD cycle (lights on/off at 07:00 and 19:00, respectively); fed three times per day with dry food (TetraMin, Tetra, Germany) mainly in the first part of the light hours (at 09:00; 11:00 and 13:00). Fish were reared and manipulated following the guidelines of the European Union (2010/63/EU) and the Spanish legislation (RD 1201/2005 and Law 32/2007). The experimental protocols were approved by the Spanish National Committee and the Committee of the University of Murcia on Ethics and Animal Welfare.

Behavioral traits and circadian rhythms are sensitive to social modulation (Castillo-Ruiz, Paul & Schwartz 2012; Bloch *et al.* 2013; Jolles *et al.* 2017), and previous results showed that the selection lines behaved differently when tested in isolation or in groups (Sbragaglia *et al.* 2019a). Indeed, being in a group has important consequences for foraging behavior in zebrafish (Pitcher 1986; Harpaz & Schneidman 2019), and social isolation can create anxiety-related behaviors (Shams *et al.* 2017). Therefore, knowing that zebrafish is a social species (Spence *et al.* 2008), and the size-selective harvesting treatments occurred in a social environment, we studied zebrafish behavior at the group level for testing our hypotheses.

### Collective risk-taking behavior experiments

We first characterized the diving behavior of zebrafish to test our first hypothesis (H1 in Fig. 1). Groups of eight juveniles (30 days post fertilization) were stocked into 3 liter rearing boxes. The boxes (N = 36) were housed on the shelves of the same zebrafish holding system with a randomized order (6 replicates for each line; 12 replicates per treatment). Throughout the experiment zebrafish were fed ad libitum with dry food (TetraMin, Tetra, Germany) and maintained in the same conditions reported above. Measurements of diving behavior were assessed at 230 and 240 days post fertilization to estimate consistent inter-group differences as a measure of collective personality. To that end, the groups of eight zebrafish were moved from the 3-liter rearing box in a new experimental tank (width x length x height = 10 x 30 x 25 cm) with 22 cm of water (Fig. S2). The experimental tank was placed on a table behind a white curtain. On the side of the experimental tank (at about 50 cm), we placed a webcam (Fig S2 and video S1) and measured the time zebrafish shoals spent at the surface during an experimental assay composed of an acclimation period and feeding under the risk of predation (Fig. S2 and Video S1). The time spent at the surface is a well-established behavioral test used in zebrafish to measure anxiety-like behavior (Levin, Bencan & Cerutti 2007; Egan *et al.* 2009; Kalueff 2017). Zebrafish show a typical diving response moving towards the bottom of the tank as soon as they are introduced in a novel tank, followed by a slow exploration of the surface (Kalueff 2017). The surface of the water is a risky environment for zebrafish (Spence *et al.* 2008), and previous experimental results showed that stimuli mimicking an approaching predator from the top of the tank induced a robust increase of time spent at the bottom of the tank (Luca & Gerlai 2012). Therefore, we expected an incremental use of the water surface during the acclimation period (just after the zebrafish are introduced in the novel experimental tank), with its maximum happening where food is added at the surface, followed by a drastic reduction after the approach of a simulated predator from the top (Fig. S2 and Video S1).

To capture the quantitative development of the behavior of the shoal throughout the experimental assay, we subdivided the observations in 30-s periods. Food was added after 3 min, and then after 30 s a simulated bird (Fig. S2 and Video S1) approached from the top of the experimental tank. The simulated bird was released with a cable from a height of 1 m (Fig. S2 and Video S1) and was maintained above the tank for 5 s. By repeating the experiment 10 days later we estimated the repeatability of the behavior of shoals.

### Circadian experimental set up and design

To test our second hypothesis (H2 in Fig. 1) we recorded daily activity rhythms in a parallel experiment with a new set of fish when they were at about 170 days post fertilization. We randomly selected 3 groups of 15 fish for each of the six selection lines (6 groups and 90 fish for each treatment: large-harvested, small-harvested and control line). The individual mass of the experimental fish groups was within the following range: large-harvested (3.6 – 5.7 g); control (3.3 – 6.5 g); small-harvested (4.6 – 6.1 g). The experimental set up was designed to jointly record swimming (as a proxy of locomotor activity) and self-feeding activity by a group of zebrafish (Fig. S3; see also del Pozo, Sanchez-Ferez & Sanchez-Vazquez 2011).

Before starting experimental trials, fish were acclimated to laboratory conditions and to the use of the self-feeder for 15 days. Afterwards we recorded the swimming and self-feeding activity for 38 days (from at about 170 to 210 days post fertilization) following stablished protocols in chronobiology to test for daily and circadian rhythmicity of fish behaviour (e.g. Sánchez-Vázquez & Tabata 1998). The experimental trial was subdivided in four consecutive phases: (first light-darkness trial: 15 days) 12:12 h light-darkness cycle to investigate daily swimming and self-feeding rhythms; (constant conditions: 7 days) constant dim light (3.5 lux at the water surface) to investigate the endogenous free-running rhythms of swimming and self-feeding activity; (second light-darkness trial: 15 days) 12:12 h light-darkness cycle to investigate their resynchronization; (final constant conditions) constant dim light to ascertain the endogenous expression patterns of clock genes.

### Gene expression analysis

We measured gene expression at the end of the activity rhythms to test our third hypothesis (H3 in Fig. 1). We selected seven genes controlling the most important functioning mechanisms of the circadian system in fish (Vatine *et al.* 2011; Idda *et al.* 2012; Foulkes *et al.* 2016; Frøland Steindal & Whitmore 2019). The first group of genes was composed by genes related to the core transcriptional-translational feedback loop driving circadian oscillation in vertebrates (*per1b, clock1a, arntl1a*). The second group involved light-inducible genes (*per2, cry1a*). The third group was composed of genes related to the circadian clock output (*dbpa, tef1*). Finally, a fourth group was composed of circadian clock controlled genes (*lepa*, and *igf1*) related to growth and energy balance. *lepa* was measured both in brain and liver and *igf1* only in liver. The product of the gene *lepa* acts as satiation signal in teleosts and has implications in energy balance and glucose homeostasis, it is mainly secreted in the liver with a daily rhythm of expression (Paredes *et al.* 2015; Rønnestad *et al.* 2017). The gene *lepa* is also expressed in the teleost brain, where expression increases after a meal (Yuan *et al.* 2016). The gene *igf1* is involved in growth processes and signals lipid metabolism and it is secreted with a daily rhythm of expression (Paredes *et al.* 2015).

Samples were collected throughout a 24-hour cycle at 6 different time points (Time of the day: 21, 01, 05, 09, 13, 17). Fish were exposed for 24 h to the second constant dim light phase. We sampled a total of 5 fish for each time point and each treatment. Zebrafish were euthanized on ice and then the whole brain and liver were dissected and put in RNAlater at −80°C. Total RNA was isolated using a Trizol reagent (Invitrogen, USA) following the manufacturer’s instructions. The amount, quality and composition of isolated RNA were analyzed by Nanodrop ND-1000 (Thermo Fisher Scientific Inc., Wilmington, USA). DNase-treated RNA was used to perform cDNA synthesis in a final volume of 20 μl using the iScript cDNA synthesis kit (Biorad, USA). The reaction was performed at 46°C for 20 min, followed by a 1-min inactivation step at 95°C.

cDNA was PCR-amplified with the StepOnePlus Real-Time PCR System (Applied Biosystems, Foster City, CA, USA) using SsoAdvanced™ Universal SYBR® Green Supermix (Biorad, USA). The thermal cycling conditions were as follows: 30 s of denaturation at 95°C, followed by 40 cycles of a 15-s denaturation step at 95°C, and then by an annealing-elongation step for 30 s at 60°C. After amplification, a melting curve analysis was performed to confirm amplicon specificity. The samples were run in triplicate. Gene-specific primers are indicated in Table S1. The relative expression levels of each sample were calculated by the 2^-ΔΔCT^ method (Livak & Schmittgen 2001), using geometric mean of three housekeeping genes (*βactin, GADPH* and *EF1α*) in both tissues.

### Statistical analysis

Risk-taking data were power transformed (more details in text S2), and adjusted repeatability (i.e., after controlling for fixed effects of selection lines; Nakagawa & Schielzeth 2010) was calculated for each 30-s time bin of the time spent at the surface across time (230-240 days post fertilization). The replicates of the selection lines were used as additional random effect into the model.

We split the results related to risk-taking in two time-bin sections: (*i*) before the approach of the simulated predator; (*ii*) after the approach of the simulated predator. In both cases we modelled surface time (square root transformed) by using non-orthogonal quadratic polynomial mixed models with selection lines as fixed factor (three levels) and consecutive 30-s time bins were used as covariate. The two lines replicates were nested within each trial and treated as random intercepts.

Waveform analysis (24-h based) was carried out in both LD trials, and raw time series values (i.e. number of infrared light beam interruptions) were transformed in % of maximum to standardize the data. The Midline Estimating Statistic Of Rhythm (MESOR) was computed and represented as a horizontal threshold in waveform plots (Refinetti 2006).

Regarding swimming activity rhythms, we estimated total activity for each group by calculating the area under the waveform curve during both LD trials (Refinetti 2006). Next, early peaks of activity were calculated as the ratio between the percentages of area under the curve divided by the percentage of time of the period considered (i.e. four hours; Sbragaglia *et al.* 2013). Regarding self-feeding activity rhythms, we estimated feeding events for each group by calculating the area under the waveform curve during four different hourly sections (i.e. where most of the activity was concentrated): *i*) from 06:00 to 07:00; *ii*) from 07:00 to 08:00; *iii*) from 08:00 to 09:00 (i.e., the hour before lights on); *iv*) from 09:00 to 10:00 (i.e., the hour after lights on). After a power transformation of the data, adjusted repeatability (selection lines as fixed effect and lines replicates as additional random effect into the model) were calculated over the two LD trials. Next, we modelled the data using a linear mixed-effects model with selection lines as fixed effect and line replicates nested within LD trials as random intercepts.

Finally, circadian periodicity in gene expression was assessed using RAIN, a robust non-parametric method for the detection of rhythms in biological data that can detect arbitrary waveforms and has been widely applied in the measurement of circadian transcripts abundance (Thaben & Westermark 2014).

Periodogram and waveform analysis were performed using the software Eltemps (www.el-temps.com). The rest of analyses were implemented using R 3.5.0 (https://www.R-project.org/) and the following additional packages; “rain” (Thaben & Westermark 2014); *“rcompanion”* package for power transformation (https://CRAN.R-proiect.org/package=rcompanion), “rptR” for calculating repeatability (Stoffel, Nakagawa & Schielzeth 2017), “lme4” R package (Bates *et al.* 2014) to implement the linear mixed models, *“MuMIn”* to get marginal and conditional R^2^ (Bartoń 2014). We used a 95% confidence interval.

## RESULTS

### Collective risk-taking behavior

Time spent at the surface was overall significantly repeatable (adjusted R ranged from 0.22 to 0.55; Table S2) - after accounting for the fixed effects of selection lines - across time (230 and 240 days post fertilization) with some differences depending on the 30-s time period considered (Table S2; Fig. 2; see text S2). We found a decrease of the adjusted repeatability towards the end of the experimental assay (Fig. 2; see Fig. S4 for variance partitioning).

**Figure 2.**
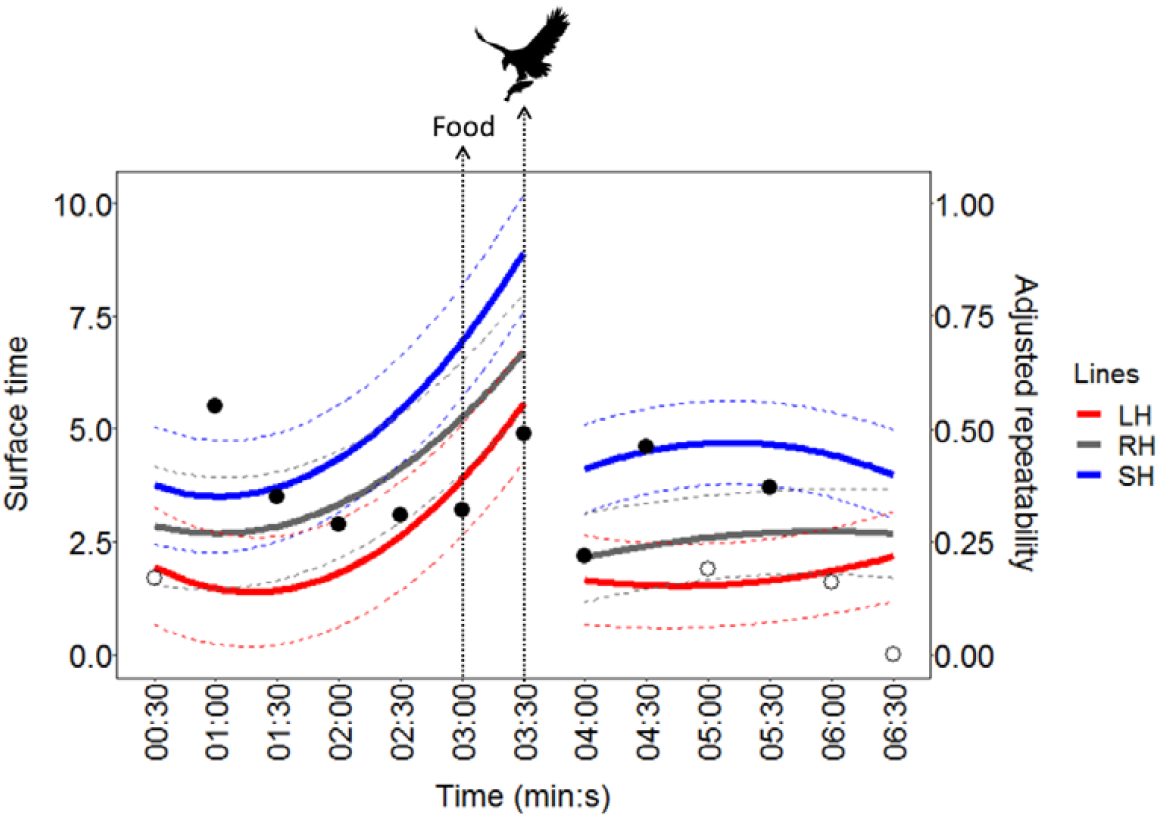
Results of the diving test used to assess collective risk-taking behavior in the zebrafish selection lines in two consecutive trials at 230 and 240 days post fertilization. Results are presented and modelled as cumulative time (bin in 30-s period) spent by zebrafish shoals at the surface of the water (7 cm; see Fig. S2 and video S1) once moved to a novel tank (left axis) and adjusted repeatability scores (right axis). Food has been added at 03:00 and the predator approached at 03:30. The colored lines represent the fit of the surface time (left axis) with a non-orthogonal quadratic polynomial (*N* = 12 for each selection line) together with confidence intervals (dashed colored lines). Repeatability scores (right axis) are presented as empty circles (when they were not significantly different from zero) and solid circles (when they were significantly different from zero; see also table S1-2 and figure S4 for partitioning of variance components). Bird pic source: https://www.piqsels.com.

The time spent at the surface before the approach of the simulated predator followed a positive quadratic trajectory, which was similarly expressed in all selection lines (Table S3; Fig. 2). All zebrafish initially avoided the surface after being introduced into the experimental tank, but soon after started to increase the time spent at the surface reaching the maximum when the food was added (03:30 in Fig. 2). After the approach of the simulated predator (04:00 in Fig. 2), all the selection lines strongly reduced their time spent at the surface (Table S3; Fig. 2). The small-harvested line consistently indicated to be more risk-taking than controls, by consistently spending more time at the surface, while the opposite was observed for the large-harvested line (Fig. 2). Therefore, we can conclude that the large-harvested line was shyer and the small-harvested line was bolder than the control.

### Behavioral activity rhythms

The behavioral rhythms of all the selection lines revealed significant 24-hour periodicities during both LD trials (first LD trial, days 1-15; second LD trial, days 22-36) and significant free-running rhythmicity under constant dim light conditions, with periods close to 24-h (Fig. S5, Table 1, see text S3). All lines displayed typical diurnal swimming activity rhythms with greater activity during photophase (Fig. 3A-B).

**Figure 3.**
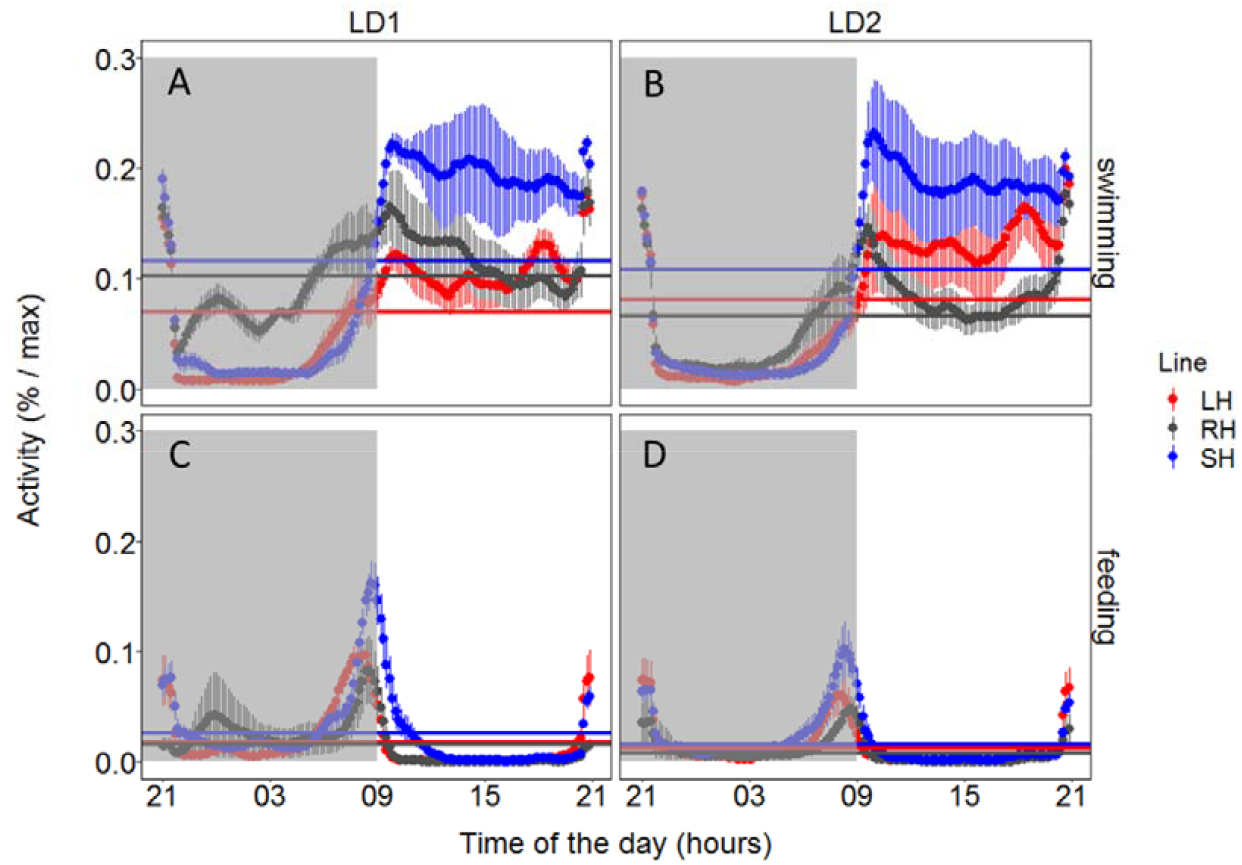
Mean waveforms (time scale 24 h) for swimming (A, B) and self-feeding (C, D) activity of zebrafish shoals are expressed as percentage of the maximum during the first (LD1: days 1-15) and second (LD2: days 22-36) light-darkness trial. Each point represents the 10-min binned mean across all the experimental days in LD1 and LD2 for all the groups. The different colors represent the selection lines: red for large-harvested (LH; N = 6) grey for control (RH; N = 6) and blue for small-harvested (SH; N = 5). The horizontal lines represent the midline estimating statistic of rhythm as reported in Table 1. The vertical lines represent the standard error (*N* between 75 and 90). Grey shadowed areas represent the darkness hours (lights on is at time of the day = 09:00).

**Table 1.** The output of the periodogram analysis for swimming and self-feeding activity rhythms during the three steps of the experiment (first light-darkness trial: LD1; constant dim light: LL1; and second light-darkness trial: LD2) according to the three selection lines (large-harvested: LH, small-harvested: SH and control: RH). Periods (T) or free running periods (*tau*) are averaged according to the number of groups (N; the proportion indicates the number of groups with significant rhythms) and the robustness of periodicities are expressed as percentage of variance explained (%V) together with the midline estimating statistic of rhythm (MESOR). The standard errors are reported between parentheses.

| Phenotype | Experimental Phase | Line | N | T or *tau* (se) | %V (se) | MESOR (se) |
| --- | --- | --- | --- | --- | --- | --- |
| Swimming | LD1 | LH | 6/6 | 24.00 (0.00) | 47.41 (4.54) | 0.070 (0.006) |
|  |  | RH | 6/6 | 24.00 (0.02) | 29.74 (10.38) | 0.103 (0.011) |
|  |  | SH | 5/6 | 24.02 (0.02) | 52.48 (10.13) | 0.116 (0.012) |
|  | LL1 | LH | 4/6 | 20.88 (1.70) | 29.43 (11.23) | - |
|  |  | RH | 2/6 | 26.33 (0.00) | 30.29 (8.59) | - |
|  |  | SH | 2/6 | 23.42 (0.33) | 23.54 (1.89) | - |
|  | LD2 | LH | 6/6 | 24.03 (0.02) | 52.45 (8.54) | 0.081 (0.017) |
|  |  | RH | 6/6 | 24.00 (0.03) | 51.01 (14.30) | 0.066 (0.006) |
|  |  | SH | 5/6 | 24.00 (0.00) | 56.96 (20.82) | 0.108 (0.018) |
| Self-feeding | LD1 | LH | 6/6 | 24.00 (0.00) | 46.44 (7.45) | 0.018 (0.003) |
|  |  | RH | 6/6 | 23.97 (0.02) | 38.89 (18.18) | 0.016 (0.009) |
|  |  | SH | 6/6 | 24.01 (0.03) | 43.21 (6.35) | 0.026 (0.005) |
|  | LL1 | LH | 4/6 | 23.56 (0.52) | 28.59 (6.73) | - |
|  |  | RH | 3/6 | 22.56 (1.31) | 18.86 (3.43) | - |
|  |  | SH | 1/6 | 16.83 (-) | 16.03 (-) | - |
|  | LD2 | LH | 6/6 | 24.03 (0.02) | 46.07 (11.67) | 0.013 (0.003) |
|  |  | RH | 6/6 | 24.01 (0.01) | 32.29 (7.92) | 0.008 (0.003) |
|  |  | SH | 6/6 | 24.00 (0.02) | 37.87 (10.17) | 0.016 (0.003) |

Total swimming activity during the photophase was significantly (*p* < 0.001) repeatable - after accounting for the fixed effects of selection lines - over the two LD trials (R = 0.78 [CI: 0.27 – 0.92]; Fig. S6A). The small-harvested was significantly (t_8_ = 3.52; *p* < 0.01) more active than controls (Fig. 4A), while no significant differences (t_8_ = 0.71; *p* = 0.495) were detected between the large-harvested line and controls (Fig. 4A). Early peaks of swimming activity were also significantly (*p* < 0.05) repeatable (R = 0.58 [CI: 0.09 – 0.83], after accounting for the fixed effects of selection lines (Fig. S6B). In particular, the small-harvested line concentrated significantly (t_8_ = 2.96; *p* < 0.05) more daily activity on the first four hours of the photophase relative to controls (Fig. 4B), while no significant differences (t_8_ = 0.61; *p* = 0.558) were detected between the large-harvested line and controls (Fig. 4B).

**Figure 4 –.**
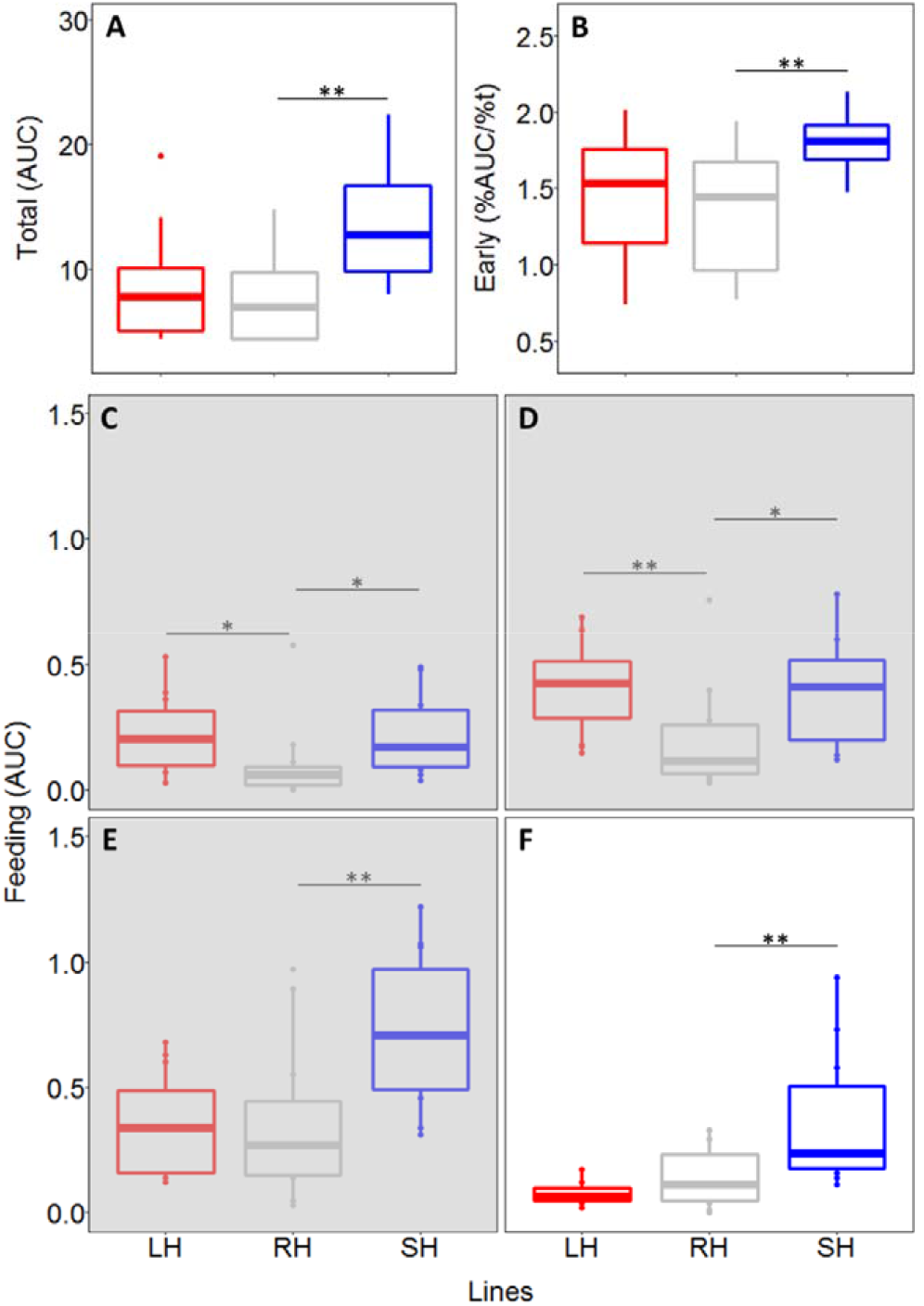
Total swimming activity during light hours (A; area under the waveform curve, AUC) and early daily activity (B; percentage of activity during the first four hours of light) together with self-feeding activity (C-F; area under the waveform curve, AUC) during the last three hours of scotophase (grey shadow; C = 06:00 - 07:00; D = 07:00 - 08:00; E = 08:00 - 09:00) and the first hour of photophase (F = 09:00 - 10:00). The different colors represent the selection lines: red for large-harvested (LH) grey for control (RH) and blue for small-harvested (SH); N between 5 and 6 (see table 1 for more details). Boxplots represent the median (bold centerline), the 25^th^ (top of the box) and 75^th^ percentile (bottom of the box). Significant differences are indicated by black horizontal lines (*: *p* value < 0.05; **: *p* value < 0.01).

The lines displayed self-feeding activity rhythms during the last hours of darkness (Fig. 3C-D). The self-feeding activity was significantly (*p* < 0.05) repeatable (adjusted R ranged from 0.45 to 0.52) over the two LD trials in the four consecutive hourly sections that were analyzed (from three hours before to one hour after light on, Fig. S6C-F). At the onset of the self-feeding events (06:00 - 07:00 h; darkness) both size-selected lines were significantly (large-harvested line: t_8_ = 2.94, *p* < 0.05; small-harvested line: t_22_ = 2.85, *p* < 0.05) more active than controls (Fig. 4C). The same pattern (large-harvested line: t_8_ = 3.37, *p* < 0.01; small-harvested line: t_8_ = 3.10, *p* < 0.05) was detected in the next hourly section (07:00 - 08:00 h; darkness, Fig. 4D). Subsequently, during the hour before light-on (08:00 - 09:00 h), the small-harvested line was significantly (t_8_ = 3.64, *p* < 0.01) more active in demanding food than controls (Fig. 4E), while the large-harvested line did not show significant differences respect to control (t_8_ = 0.61, *p* = 0.558). The same pattern (large-harvested line: t_8_ = −0.43, *p* = 0.680; small-harvested line: t_8_ = 3.49, *p* < 0.01) was detected during the first hour after light-on (09:00 −10:00 h; Fig. 4F).

### Circadian rhythms in gene expression

Core clock genes expression revealed significant differences in their circadian oscillations among the zebrafish lines (Table 2). In the brain, the control line showed a strong trend (*p* = 0.052) in the circadian rhythmicity of *arntl1a* and a clear circadian oscillation of *clock1a* with a peak at the same time of the day (TD) of *arntl1a* (TD09 for both genes; Table 2). By contrast, the small-harvested line did not reveal circadian rhythmicity of *arntl1a* and *clock1a* compared to the control, while the large-harvested line shifted its peaks of circadian expression for both genes by 8 hours (TD01 instead of TD09, Table 2 and Fig. 5). Results of *per1b* indicated significant circadian patterns in the three lines with main differences in peaks in both size-selected lines that shifted from TD17 to TD21 relative to the control (Fig. 5, Table 2). In the liver, the small-harvested line showed similar circadian rhythmicity to the control line in terms of *arntl1a*, while the large-harvested line revealed only a trend (*p* = 0.086) in the circadian oscillation (Table 2). The control line did not show any circadian rhythmicity in terms of *clock1a* and *per1b*, however, the size-selected lines revealed clear circadian rhythmicity for these two genes with slightly different peaks for *per1b* (Table 2 and Fig. 5).

**Figure 5.**
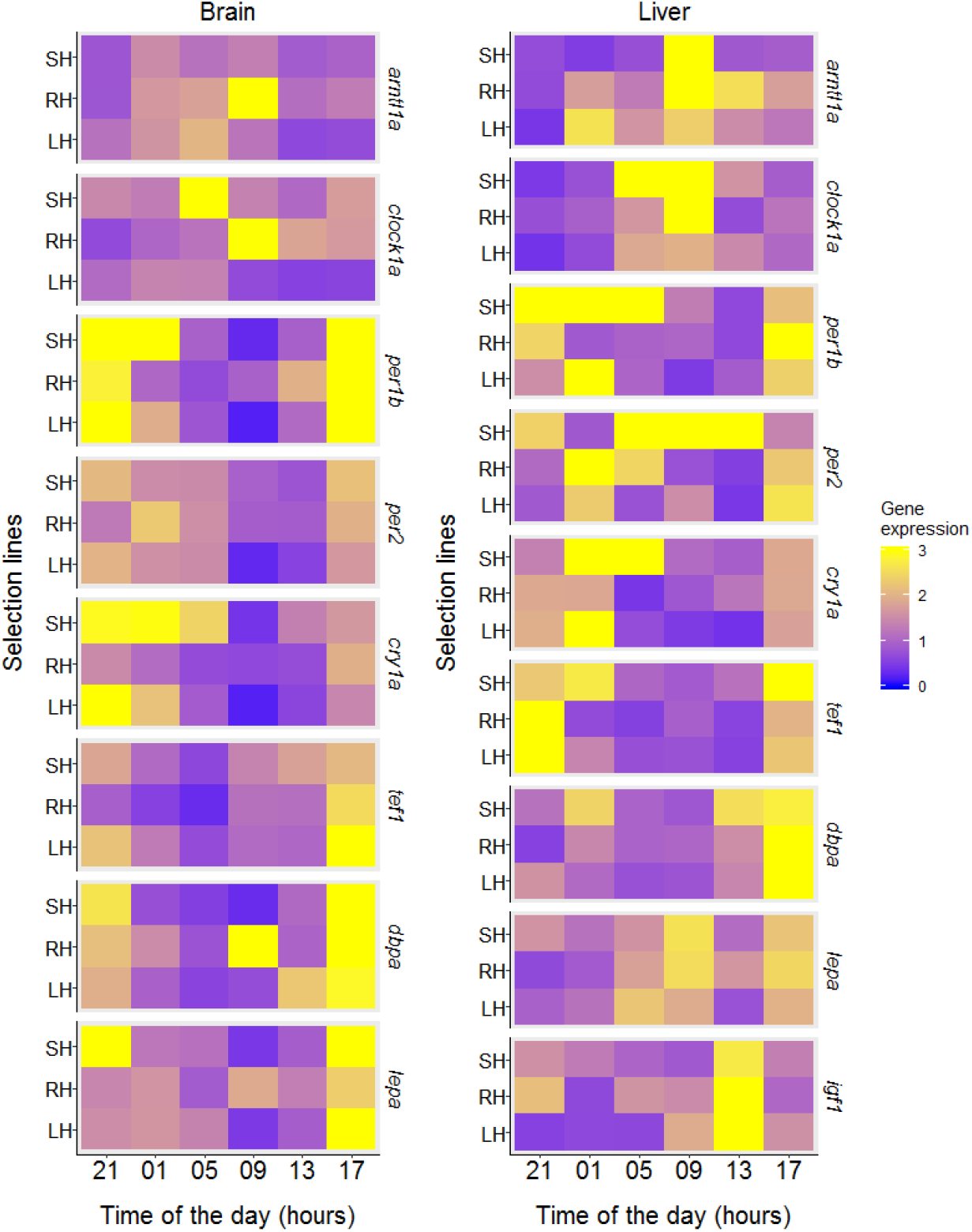
Mean endogenous (i.e. measured under constant dim light) transcript abundance of the genes related to the circadian system and clock controlled genes related to growth and energy balance in the brain (left) and liver (right) at different time of the day for the three selection lines (large-harvested: LH, small-harvested: SH and control: RH; N between 4 and 5, see table 2 for more details).

**Table 2.** The output of the analysis with RAIN (a robust non-parametric method for the detection of rhythms in biological data) regarding the circadian oscillation of transcript abundance of the genes related to the circadian system to neuroendocrinological aspects of energy balance, appetite and growth. Gene expression was measured at the end of the experiment in constant dim light. Transcripts are subdivided according to their role for each selection lines (large-harvested: LH, small-harvested: SH and control: RH) and in each tissue (brain and liver). The timing of the peak of the circadian oscillation is indicated using the time of the day in hours (see also Fig. 5) together with the *p* value and total N (N is between 4 and 5 for each time point and selection line). Significant circadian rhythmicity is reported in bold.

| Clock mechanism | Gene | Line | Brain |  |  | Liver |  |  |
| --- | --- | --- | --- | --- | --- | --- | --- | --- |
|  |  |  | N | Peak | *p* value | N | Peak | *p* value |
| Core clock | *arntl1a* | LH | **29** | **01** | **<0.01** | 30 | 01 | 0.086 |
|  |  | RH | 26 | 09 | 0.052 | **30** | **09** | **<0.001** |
|  |  | SH | 30 | 01 | 0.682 | **30** | **09** | **<0.01** |
|  | *clock1a* | LH | **28** | **01** | **<0.01** | **28** | **09** | **<0.001** |
|  |  | RH | **28** | **09** | **<0.05** | 30 | 09 | 0.181 |
|  |  | SH | 30 | 05 | 0.279 | **30** | **09** | **<0.001** |
|  | *per1b* | LH | **29** | **21** | **<0.001** | **29** | **01** | **<0.01** |
|  |  | RH | **29** | **17** | **<0.01** | 30 | 17 | 0.751 |
|  |  | SH | **29** | **21** | **<0.001** | **29** | **21** | **<0.05** |
| Light-inducible | *per2* | LH | **30** | **21** | **<0.001** | 30 | 01 | 0.376 |
|  |  | RH | 27 | 01 | 0.117 | 30 | 01 | 0.054 |
|  |  | SH | **28** | **17** | **<0.05** | 30 | 05 | 0.308 |
|  | *cry1a* | LH | **30** | **21** | **<0.001** | **28** | **21** | **<0.01** |
|  |  | RH | **30** | **17** | **<0.001** | **30** | **01** | **<0.01** |
|  |  | SH | **30** | **01** | **<0.001** | 30 | 05 | 0.120 |
| Clock output | *tef1* | LH | **29** | **17** | **<0.05** | **30** | **17** | **<0.05** |
|  |  | RH | **30** | **17** | **<0.001** | **30** | **21** | **<0.001** |
|  |  | SH | **29** | **17** | **<0.05** | **30** | **17** | **<0.05** |
|  | *dbpa* | LH | 29 | 17 | 0.078 | 30 | 17 | 0.144 |
|  |  | RH | **30** | **17** | **<0.05** | 29 | 17 | 0.205 |
|  |  | SH | **29** | **17** | **<0.001** | 28 | 17 | 0.190 |
| Appetite<br>Food intake | *lepa* | LH | **27** | **17** | **<0.05** | 27 | 05 | 0.769 |
|  |  | RH | 30 | 17 | 0.456 | 28 | 17 | 0.319 |
|  |  | SH | **30** | **17** | **<0.01** | 30 | 17 | 0.876 |
| Growth | *lgf1* | LH | - | - | - | 28 | 13 | 0.248 |
|  |  | RH | - | - | - | **28** | **13** | **<0.001** |
|  |  | SH | - | - | - | 30 | 13 | 0.134 |

### Light-inducible genes

In the brain, the control line showed no significant rhythmicity of *per2*, while the size-selected lines oscillated with peaks at different times (TD21 and TD17 for large- and small-harvested lines, respectively; Table 2, Fig. 5). Results of *cry1a* indicated significant circadian rhythmicity with different peaks in the size-selected lines relative to the control: the large-harvested line shifted the peak from TD17 to TD21, while the small-harvested line shifted from TD17 to TD01 (Table 2 and Fig. 5). In the liver, both size-selected lines lost circadian rhythmicity of *per2* with respect to the control that showed a strong trend (*p* = 0.054). The small-harvested line lost rhythmicity for *cry1a*, while the peak of the large-harvested line shifted from TD01 to TD21 compared to the control (Table 2 and Fig. 5).

### Clock output genes

The circadian expression of clock output genes showed less peak shifts compared to the other two groups of genes that we investigated (Table 2). In the brain, the three lines had a circadian rhythmicity of *tef1* and *dbpa* with peaks at the same time (TD17). However, the large-harvested line showed a trend (p = 0.078) in the circadian rhythmicity of *dbpa* (Table 2 and Fig. 5). In the liver, both size-selected lines shifted the peak of the circadian oscillation of *tef1* from TD21 to TD17 compared to the control (Table 2 and Fig. 5). By contrast, the three lines lost circadian rhythmicity in terms of *dbpa* (Table 2 and Fig. 5).

### Clock controlled genes related to growth and energy balance

The circadian expression of genes related to physiological processes, such as energy balance (*lepa*) and growth (*igf1*), also revealed significant differences between the size-selected lines and controls. The circadian oscillation of *lepa* expression in the brain was only significant in the size-selected lines, but not in the control, with peaks at the same time (TD17; Table 2 and Fig. 5). By contrast, we did not find significant circadian oscillation of *lepa* in the liver (Table 2, Fig. 5). Finally, the *igf1* circadian oscillation in the liver was significant only in the control line (Table 2, Fig. 5).

## DISCUSSION

We found that five generations of size-selective harvesting left an evolutionary legacy in the collective risk-taking behavior as well as in the circadian behavioral and molecular outputs. Three key results are noteworthy and further discussed below. First, selective harvesting of larger individuals fostered evolutionary changes of collective risk-taking behavior. Thereby, we provide experimental evidence to theoretical predictions that a fast-life history can coevolve with a reduced risk-taking behavior (Andersen, Marty & Arlinghaus 2018; Claireaux, Jorgensen & Enberg 2018), in contrast to predictions of the pace-of-life syndrome (Réale *et al.* 2010). Second, we found risk-taking behavior and early daily locomotor activity to be linked only in the small-harvested line. By contrast, the large-harvested line did not show differences in early locomotor activity compared to the control, suggesting asymmetric evolutionary changes induced by size-selective harvest in relation to the fine-scale timing of swimming and self-feeding activity. Finally, from a molecular perspective, our data suggest the presence of a switch in the circadian output pathway that substantially reverted the molecular circadian clockwork (Fig. 6). In fact, we found strong changes in the molecular circadian clockwork of both size-selected lines, but limited changes at the behavioural level, e.g. all lines were clearly diurnal. We suggest being active during light hours and look for food during the last hours of darkness may have important adaptive value for zebrafish, no matter what the molecular clock is signaling. Therefore, our results suggest a strong evolutionary resistance (*sensu* Sgrò, Lowe & Hoffmann 2011) of daily behavioral rhythms in response to size-selective harvesting.

**Figure 6.**
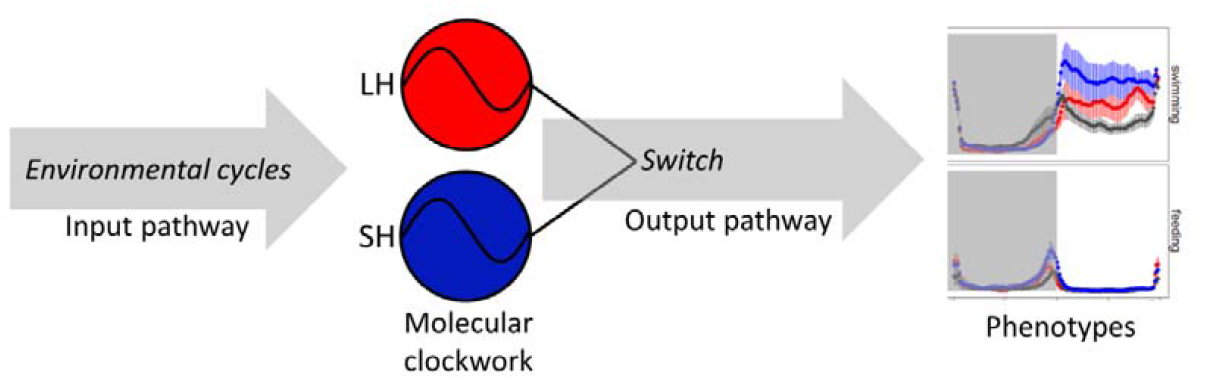
The conceptual framework of the downstream switch we proposed to explain the results presented here. The circadian clock synchronized with the environmental cycles (e.g. light-darkness or feeding). Next, the size-selective harvesting (large-harvested in red: LH; small-harvested in blue: SH) triggers the evolution of different molecular circadian clockwork with respect to control. Finally, in the output pathway the different molecular circadian clockwork are switched to produce a similar output driving the daily activity rhythms phenotypes (i.e. swimming and self-feeding activity rhythms).

### Collective risk-taking behavior

We found collective foraging in a risky context to be a repeatable behavior and thus indicative of zebrafish collective personality traits (Bengston & Jandt 2014; Jolles *et al.* 2018). We also documented that consistency of collective behavior can vary during an experimental assay as demonstrated before for individual traits (e.g. O’Neill *et al.* 2018). We found support for our first hypothesis (H1) in terms of the large-harvested line decreasing collective risk-taking behavior (and hence becoming shyer), while the small-harvested line increasing collective risk taking (and hence becoming bolder) compared to the control. The match/mismatch between the species-specific evolutionary history and human-induced rapid environmental changes is key to understand behavioral responses, such those observed in our experimental system (Sih, Ferrari & Harris 2011). The surface of the water is a risky environment for zebrafish (Spence *et al.* 2008; Cachat *et al.* 2011), therefore it is expected that the time spent feeding at the surface is under predation-driven selection pressure in wild populations and that genetic variation for surface feeding is present in the wild (Sih, Ferrari & Harris 2011). We started our experimental system with wild zebrafish populations and fed them with clumped food at the surface (Uusi-Heikkilä *et al.* 2015); therefore the time spent feeding at the surface played a major role in determining size-at-age – the trait we selected on. Our selection pressure was divergent, either favoring large or small size at-age, which could explain the symmetrical evolutionary changes we documented in the two opposing selection lines in terms of collective risk-taking behavior. Our interpretation is that our selection pressure favored fish that were either willing (small-harvested line) or not willing (large-harvested line) to take risk during foraging and therefore behavioural changes in terms of collective risk-taking behavior followed the selection on size-at-age through a correlated selection response. Such interpretation would imply that the evolutionary changes of risk-taking behavior were driven by an energy acquisition pathway (Enberg et al. 2012), coupling behavior and size-at-age such that selection on size-at-age also alters risk-taking behavior (Biro & Post 2008). An alternative explanation in terms of life-history adaptations fostering characteristic changes in risk-taking behavior (e.g. evolution of a fast life-history correlating with increased boldness; Claireaux, Jorgensen & Enberg 2018) are less supported by our data. In fact our results did not show an increase of risk-taking behavior in the large-harvested line as expected according to their fast life history (Uusi-Heikkilä *et al.* 2015). We found the opposite.

Previous work in our laboratory using the same experimental system was conducted at the individual level, and results related to individual risk-taking behavior in response to size-selection revealed more ambiguous patterns than those reported here at the collective level. Specifically, Uusi-Heikkilä *et al.* (2015) reported that juveniles of the small-harvested line increased individual risk-taking compared to controls, while Sbragaglia *et al.* (2019a) showed that adult females decreased risk-taking behavior relative to controls. In these studies, however, the large-harvested line did not show changes in individual risk-taking behavior compared to controls in both juveniles (Uusi-Heikkilä *et al.* 2015) and adults (Sbragaglia *et al.* 2019a). By contrast, the present research measuring the collective risk-taking behavior showed clear symmetrical response of the size-selected lines.

Two main limitations of the previous experimental approaches that were overcome in the present research may explain the discrepancies. The first is related to the behavioral assay. Our previous work focused on individual behavioural traits using the open field test to assess risk-taking behavior (Uusi-Heikkilä *et al.* 2015; Sbragaglia *et al.* 2019a). The open field test is a commonly used behavioral assay to measure novelty-evoked and anxiety-like behaviors in fish in two dimensional horizontal space (Burns 2008; Kalueff 2017). However, it disregards the third vertical dimension, which constitutes an important behavioral aspect for zebrafish both due to its evolutionary history and because of the selection environment experienced in our experimental system where fish were held in large tanks and food was provided on the surface.

Thus, the experimental approach used in the present paper is more appropriate than the open field to reveal evolutionary changes in risk-taking behavior of the zebrafish size-selected lines. The second limitation is that our previous behavioral assays focused on isolated individuals instead of examining the behavioural phenotype of groups (Sbragaglia *et al.* 2019a). Being in a group has important consequences for foraging behavior in zebrafish (Pitcher 1986; Harpaz & Schneidman 2019), and social isolation can create anxiety-related behaviors (Shams *et al.* 2017). We propose that zebrafish expressed a more reliable behavior in the group context presented here than in the open field experiments previously reported on isolated individuals. Collectively, the evolutionary changes of risk-taking presented in the present paper seem more robust and they suggest that testing evolution of behavior in response to experimental size-selective harvest should carefully consider the match between evolutionary history of the species, the environmental conditions experienced during the artificial selection and the behavioral assay used to reveal the evolutionary basis of behavioral changes (Klefoth *et al.* 2012).

### Daily behavioral activity rhythms

We found partial support for our second hypothesis (H2) because the small-harvested line was found to be more active during the photophase and concentrated its swimming activity at the beginning of the scotophase. However, we did not find significant changes in the large-harvested line compared to controls. The changes in daily activity rhythms of the small-harvested line agree with a study demonstrating that proactive zebrafish (i.e., risk-taking individuals) are more active in the first hours of photophase compared to reactive ones (Tudorache *et al.* 2018). Further support for a link between risk-taking behavior and daily activity rhythms has been reported in a recent study that revealed artificial light at night affects risk-taking behavior in the Trinidad guppy, *Poecilia reticulata* (Kurvers *et al.* 2018). Relatedly, the self-feeding activity events of the small-harvested line extended to the hours before and after light-on. The timing of feeding is a key aspect for survival and is traded-off against predation risk and food availability (Kronfeld-Schor & Dayan 2003). The ecological significance of zebrafish feeding during the last hours of darkness may be related to avoidance of visual predators (e.g. birds; Spence *et al.* 2008) and/or to storing energy to be prepared for spawning that occurs at the beginning of photophase (del Pozo, Sanchez-Ferez & Sanchez-Vazquez 2011). However, the food demand sensor was located close to the water surface (del Pozo, Sanchez-Ferez & Sanchez-Vazquez 2011; see also Fig. S3) and an increase of self-feeding events towards light-on by the small-harvested line could also indicate additional support for the evolutionary change of collective foraging in a risky context following a resource acquisition mechanism that fostered a correlated selection response of size-at-age and risk-taking behavior.

Overall, the results related to daily behavioral rhythms revealed that the size-selected lines responded differently, indicating asymmetrical evolutionary changes in response to size selection, as previously reported for life history traits in other selection experiments with fish (Amaral & Johnston 2012; van Wijk *et al.* 2013; Renneville *et al.* 2020). Our results suggested an evolutionary link between risk-taking behavior and daily activity rhythms in the small-, but not in the large-, harvested line. Harvesting-induced evolution of behavior likely has a complex and multivariate nature (Claireaux, Jorgensen & Enberg 2018; Renneville *et al.* 2020), thereby complicating generalizations related to evolutionary correlations among traits. The somewhat asymmetrical evolutionary changes of daily activity rhythms in the two size-selected lines might be related to genetic trade-offs or other functional limitations in our model species (Arnold 1992; Renneville *et al.* 2020).

### Circadian system switch

We rejected our third hypothesis (H3) because we did not find a decrease of clock gene circadian rhythmicity (a decrease of robustness and amplitude as documented by Tudorache *et al.* 2018) linked to a decrease of collective risk-taking behavior. A possible explanation could be related to the fact that the two recent studies on zebrafish that demonstrated a link between individual personality trait and the circadian system (i.e. proactivity/reactivity; Rey *et al.* 2013; Tudorache *et al.* 2018), measured transcripts abundance under light-darkness conditions. Thereby, the truly endogenous nature of the circadian molecular clockwork could have been masked by the light-darkness cycle (i.e. direct response without synchronizing the circadian molecular clockwork; Mrosovsky 1999; Dunlap, Loros & DeCoursey 2004). In our study, we measured transcript oscillations under constant conditions, thereby providing a truly endogenous effect of size-selective harvesting on the molecular circadian clockwork.

All genes except *tef1* and *dbpa* had different peaks in the three lines, while in few cases the two size-selected lines peaked at the same time (e.g. *per1b* in the brain). Collectively, these results indicate strong differences in the circadian molecular clockwork of the size-selected lines, suggesting size-selection can alter the molecular basis of the circadian clock. However, the similar peaks recorded in the genes related to the clock output pathway (*tef1* and *dbpa*) implicated the occurrence of a switch (a plastic response in terms of distribution of activity throught the time of the day; see review by Mrosovsky & Hattar 2005) downstream to the circadian clock, especially in the brain. There are three mechanisms used to explain circadian system switches (Hut *et al.* 2012): *i*) altered properties of the circadian clock leading to different activity patterns; *ii*) identical properties of the circadian clock switching in the output pathway, leading to different activity patterns; and *iii*) identical properties of the circadian clock subjected to masking that ultimately determine the activity patterns. However, none of them fit our results. We instead suggest the presence of a fourth mechanism where altered properties of the circadian clock are switched in the molecular output pathway, leading to overall similar daily activity patterns at the behavioural level (in our case, consistency in overall diurnal activity in all zebrafish selection lines, Fig. 6). The role of the clock output in determining diurnal and nocturnal phenotypes is also supported by a recent study with the fruit fly *Drosophila melanogaster* (Pegoraro *et al.* 2018), where ten generations of artificial selection for diurnal and nocturnal phenotypes were sufficient to obtain highly divergent strains where most differentially expressed genes were associated with the clock output pathway.

The occurrence of a switch in the circadian system was also reinforced by the fact that the size-selected lines at F_9_ showed genetic changes in areas of genes associated with serotonin synthesis (Uusi-Heikkilä *et al.* 2015). Serotonin plays an important role in feeding and other behaviors in fish (Lillesaar 2011; Piccinetti *et al.* 2013; Paredes *et al.* 2015), and it is also a key element in the synthesis of melatonin (Lima-Cabello *et al.* 2014), which represents a major rhythmic output of the fish circadian system controlling photoperiodic-dependent functions and synchronization of biological processes (Falcón *et al.* 2010; Lima-Cabello *et al.* 2014). Thus, the serotonergic system could have played a role in the switching mechanism we document in response to size-selective harvest (Uusi-Heikkilä *et al.* 2015).

### Energy balance and growth

The *lepa* expression in the brain displayed significant circadian variations in both size-selected lines with a peak at the same circadian time compared with the absence of rhythmicity in the control group. Moreover, both size-selected lines showed a disruption of the *igf1* circadian rhythm in the liver. Results on both genes suggest a modification of growth, lipid and protein metabolism and energy balance (Piccinetti *et al.* 2013). Modifications of the insulin growth factors pathways in the zebrafish’s skeletal muscle have also been documented in previous size-selection experiments with zebrafish in other labs (Amaral & Johnston 2012), supporting the idea that this pathway might have a major role in mediating growth processes in response to size-selection.

### Limitations and translation of results

Although our common-garden approach controlled for much of possible non-genetic factors, subtle differences in the environment among the selection lines could have arisen from body size differences, thereby creating differences in rearing biomass density or dominance hierarchies; these environmental effects could have also shaped the collective behavioral phenotypes independently of a genetic-based legacy left by size-selective harvesting (e.g. Magnhagen 2015). However, the evolved differences in body size among selection lines were not related to the behavioural phenotypes (see text S1), supporting the idea of line-specific evolutionary responses. Nevertheless, the translation of our findings to real fisheries or natural predation contexts should be done with great care giving the experimental nature of our system. For example, in most eco-evolutionary contexts fish have multiple spawning events with overlapping generations and behavior is probably under direct harvest selection as well (Arlinghaus *et al.* 2017). Our work should be considered as experimental evidence that mortality-induced evolution of collective personality traits and the circadian system is plausible (either in response to human or non-human predators), motivating more research in the wild. Indeed, a recent study indicated that genomic basis for growth rate divergence in response to experimental size-selective harvesting recapitulated responses to size-selection gradients seen in the wild (Therkildsen *et al.* 2019). Therefore, experimental systems such as the one used here carry basic scientific information that harvesting or other forms of mortality can have evolutionary effects that extend beyond environmental effects.

## Conclusions

The main impacts of human activities on biological rhythms of wildlife are related to diurnal disturbance as documented by an increase of nocturnality in mammalian species in response to human presence (Gaynor *et al.* 2018). Another important human-induced change of activity rhythms is represented by artificial light at night that has been demonstrated to have a profound impact on circadian rhythms in a wide range of taxa (Gaston, Visser & Hölker 2015). Our work adds to this literature by showing that size-selective mortality (simulating eco-evolutionary contexts typical of fisheries or natural predation) can evolutionary change collective risk-taking behavior and consequently the circadian molecular clockwork, yet without strong changes in the daily behavioral activity rhythms of our model species. If our results hold in the wild, our study indicates that large-size selective mortality decrease risk taking behavior with likely negative consequences on the recovery of catchability after harvesting halted, while the opposite is true for small-size selective mortality. Further research needs to clarify the possibly adaptive and maladaptive consequences of harvesting-induced evolutionary changes of collective risk-taking behavior (Sbragaglia *et al.* 2019c), as well as the fitness costs of the molecular circadian switch documented here.

## Acknowledgments

We are grateful to Silva Uusi-Heikkilä for her contribution in the establishment of the selection lines (F_1_ – F_9_) and for sharing unpublished data, Andrew E. Honsey for modelling the Lester biphasic growth trajectories at F_13_ the various technicians, in particular David Lewis, for keeping the animals and Juan Fernando Paredes and José Antonio Oliver for helping during sampling for molecular analysis. We are also grateful to Horacio de la Iglesia, Achim Kramer and Carl H. Johnson for scientific discussion on preliminary results of the experiments presented here, and to reviewers for their valuable feedbacks and suggestions. V.S. was supported by a Leibniz-DAAD postdoctoral research fellowship (n. 91632699), and he is now supported by a *“Juan de la Cierva Incorporación”* research fellowship (IJC2018-035389-I) granted by the Spanish Ministry of Science and Innovation. C.B. was supported by University of Ferrara research grants (FAR2018-2019). J.F.L. was supported by project CHRONOLIPOFISH (RTI2018-100678-A-I00) and a “Ramón y Cajal” research fellowship (RYC-2016-20959) granted by the Spanish Ministry of Science and Innovation. The authors declare no conflict of interest.

## Author contributions

V.S., J.F.L., C.B. and R.A. designed research; V.S., J.F.L. and E.F. performed research; V.S. analyzed data; V.S., J.F.L., E.F., C.B., and R.A. interpreted results; V.S. and R.A wrote the paper with contributions from all the co-authors.

## Data accessibility

Data will be available on Dryad (http://datadryad.org/) upon acceptance of the paper.

## SUPPORTING INFORMATION

### SUPPORTING TEXT

#### 1. Standard length

Standard length of the fish was measured on the individuals used in the diving test experimental assay at 170 days post fertilization. The standard length of the large harvested line (2.23 ± 0.16 cm) was significantly (t_22_ = −2.30; *p* < 0.05) smaller than controls (2.35 ± 0.08 cm), while no significant differences (t_22_ = 1.24; *p* = 0.229) were detected between the small-harvested line (2.40 ± 0.08) and controls (Fig. S1). Such results agree to what was previously found by Uusi-Heikkilä *et al.* (2015) as shown in Figure S1. The same pattern was revealed at F_13_, suggesting evolutionary fixation and lack of phenotypic rebound eight generations after selection halted. Moreover, the evolved differences in standard length among selection lines were not related to the behavioural phenotypes. In fact, we never found behavioral differences of the large-harvested line respect to control as observed or standard length; conversely we have found the opposite (i.e. behavioral differences of the small-harvested line respect to control). As such, our results suggest that the collective behavioral changes are likely not associated with differences in size as previously demonstrated for individual behavioral traits (Sbragaglia *et al.* 2019a).

#### 2. Collective risk-taking behavior

Although time spent at the surface of the water are count data, the range of values was overall high (0-192 s) and a power transformation can be used as alternative approach to stabilize variance with negligible effect on the parameter estimates (O’hara & Kotze 2010). Therefore, we transformed the data by finding the exponent (lambda) which makes the values of the response variable as normally distributed as possible with a simple power transformation.

The highest repeatability scores were recorded in three different moments of the experimental trials: (*i*) just after the introduction in the new experimental set-up when zebrafish spent few time at the surface (from 00:30 to 01:30; Table S1 and Fig. 2); (*ii*) when zebrafish spent most of the time at the surface and when food was added (from 02:30 to 03:30; Table S1 and Fig. 2); (*iii*) the time period after the approach of the simulated predator (from 04:00 to 04:30; Table S1 and Fig. 2). By contrast, repeatability scores were low or not significant at the beginning and towards the end of the experimental trial (Table 1; Fig. 2). We also reported the among- and within-group variances in order to estimate which variance component contributed in changes repeatability estimates (Fig. S4). Both variances components increased at the beginning of the experimental assay and then dropped down when food was added (03:30 in figure S4), however within-group variance showed a more drastic drop respect to among-group variance. After the approach of the predator (04:00 in figure S4), the within-group variance maintained constant values, while the among-group variance continued to steadily drop towards zero (Fig. S4).

#### 3. Swimming and feeding activity rhythms

Time series of swimming and self-feeding activity were visually inspected by plotting actograms and then using a Chi-square periodogram (Sokolove & Bushell 1978) to scan for the presence of significant (*p* < 0.05) periodicity in the range 8-28 h and percentage of variance (%V) explained by each period, reported as a measure of rhythms’ robustness (Refinetti 2006). Visual inspection of actograms evidenced diurnal swimming activity (with some evident bimodal peaks at lights on/off) while feeding activity indicated a main peak in the last part of the darkness hours (Fig. S5).

The Chi-square periodogram analysis identified a significant 24-hours periodicity in swimming activity during both light-dark (LD) trials. Interestingly during the first LD trial the robustness of rhythms of the control line is lower (29.74±10.38 %V; Table 1) than the other two lines (large-harvested: 47.41±4.54 %V; small-harvested: 52.48±10.13 %V, Table 1) in fact the control line showed a higher swimming activity during darkness (Fig. 3). Such differences were no more evident during the second LD trial when all lines have similar robust rhythms (large-harvested: 52.45±8.54 %V; control: 51.01±14.30 %V; small-harvested: 56.96±20.82 %V; Table 1) and the control line did not show higher swimming activity during darkness (Fig. 3). Finally swimming activity showed variable significant free running rhythmicity under constant dim light (Table 1). The Chi-square periodogram also detected bimodality (peaks of significant 12-hours periodicity) in swimming activity in both LD trials.

Self-feeding activity rhythms had a significant 24-hours periodicity during both LD trials. Robustness was similar in all the lines during the first (large-harvested: 46.44±7.45 %V; control: 38.89±18.18 %V; small-harvested: 43.21±6.35 %V, Table 1) and second (large-harvested: 46.07±11.67 %V; control: 32.29±7.92 %V; small-harvested: 37.87±10.17 %V) LD trial. Finally under constant dim light the self-feeding activity indicated variable free running rhythmicity (Table 1). The Chi-square periodogram did not detect bimodality in self-feeding activity rhythms.

During the first LD trial bimodality was detected in 3 out of 6 groups in the large-harvested line (robustness: 25.17±12.20 %V), in three out of six groups in the control line (robustness: 8.50±1.73 %V) and in one out of five groups in small-harvested line (robustness: 23.18 %V). During the second LD trial bimodality was still present in two out of six groups in large-harvested line (robustness: 25.97±7.98 %V), in four out of six groups in the control line (robustness: 21.79±3.82), and in two out of five groups in small-harvested line (robustness: 9.17±0.97 %V).

### SUPPORTING TABLES

**Table S1 –.**
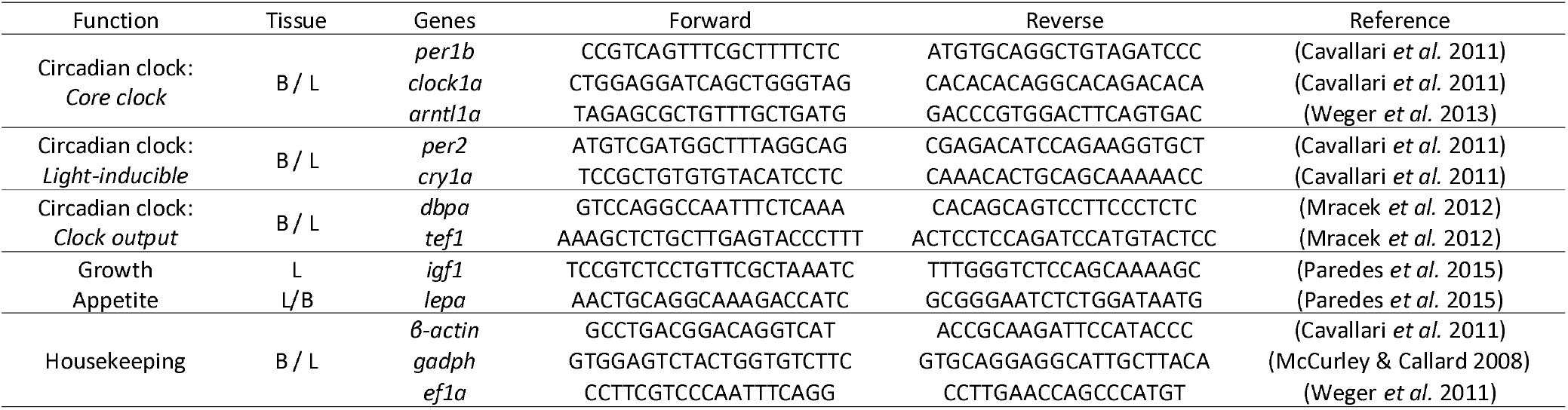
Primers used for RT-qPCR together with the references used to design them. The function of each gene is reported together with the tissue (Brain: B; Liver: L) and the forward and reverse primers.

**Table S2 –.**
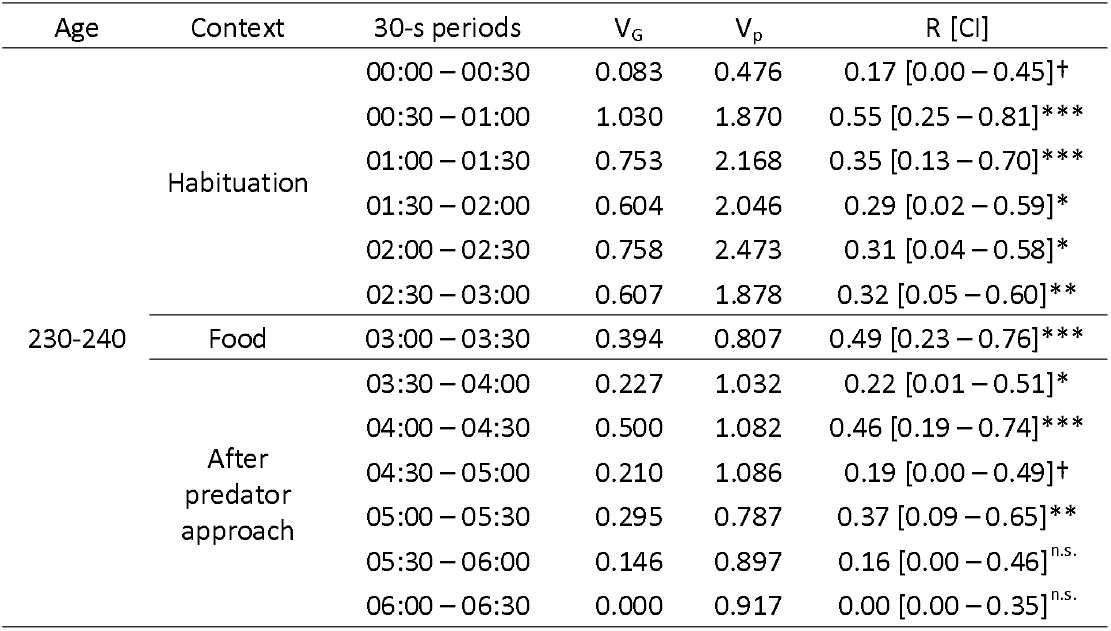
Variance partitioning (V_g_ = among-group variance; V_p_ = total phenotypic variance conditional on fixed effects; see also Fig. S4) after linear mixed-effects models. Adjusted (selection lines as fixed effect) repeatability estimates are shown together with confidence intervals (R [CI]) between two consecutive trials at 230 and 240 days post fertilization (*N* = 36 for each trial). Repeatability has been calculated for different 30-s time periods (n.s.: not significant; *: *p* value < 0.05; **: *p* value < 0.01; ***: *p* value < 0.001).

**Table S3 –.** Model estimates together with confidence intervals (lower-CI and upper-CI) and *p* values for the non-orthogonal quadratic polynomial mixed model regarding bottom dwell time measured before (from 00:00 to 03:30; including the time period where food was added) and after the approach of the predator (from 03:30 to 06:30). The 30-s time periods were used as a covariate (Time). Fish groups were nested within each line replicate and treated as random intercepts. Estimates for large-harvested (LH) and small-harvested (SH) lines are shown with respect to the control line that was randomly selected for size. The table also shows the marginal (R_m_^2^) and conditional (R_c_^2^) R-squared (*N* = 12 for each selection line).

| Response Variable | Model | Estimate | Lower-CI | Upper -CI | *p* value |
| --- | --- | --- | --- | --- | --- |
| Surface time before predator | Intercept | 3.361 | 1.829 | 4.892 | < 0.001 |
|  | Time | -0.657 | -1.323 | 0.009 | 0.055 |
|  | Time<sup>2</sup> | 0.162 | 0.081 | 0.243 | < 0.001 |
|  | Line LH | -0.486 | -2.652 | 1.680 | 0.678 |
|  | Line SH | 1.079 | -1.088 | 3.245 | 0.360 |
|  | Time × Line LH | -0.476 | -1.418 | 0.467 | 0.325 |
|  | Time<sup>2</sup> × Line LH | 0.054 | -0.061 | 0.169 | 0.357 |
|  | Time × Line SH | -0.247 | -1.190 | 0.695 | 0.609 |
|  | Time<sup>2</sup> × Line SH | 0.058 | -0.057 | 0.173 | 0.328 |
| | $R_m^2 = 0.32$ ; $R_c^2 = 0.68$ | | | | |
| Surface time after predator | Intercept | 1.811 | 0.665 | 2.957 | < 0.05 |
|  | Time | 0.377 | -0.183 | 0.938 | 0.190 |
|  | Time<sup>2</sup> | -0.038 | -0.117 | 0.040 | 0.340 |
|  | Line LH | 0.063 | -1.554 | 1.680 | 0.942 |
|  | Line SH | 1.704 | 0.089 | 3.321 | 0.057 |
|  | Time × Line LH | -0.649 | -1.441 | 0.144 | 0.111 |
|  | Time<sup>2</sup> × Line LH | 0.092 | -0.019 | 0.203 | 0.107 |
|  | Time × Line SH | 0.324 | -0.466 | 1.114 | 0.425 |
|  | Time<sup>2</sup> × Line SH | -0.065 | -0.176 | 0.045 | 0.252 |
| | $R_m^2 = 0.25$ ; $R_c^2 = 0.72$ | | | | |

### SUPPORTING FIGURES

**Figure S1.**
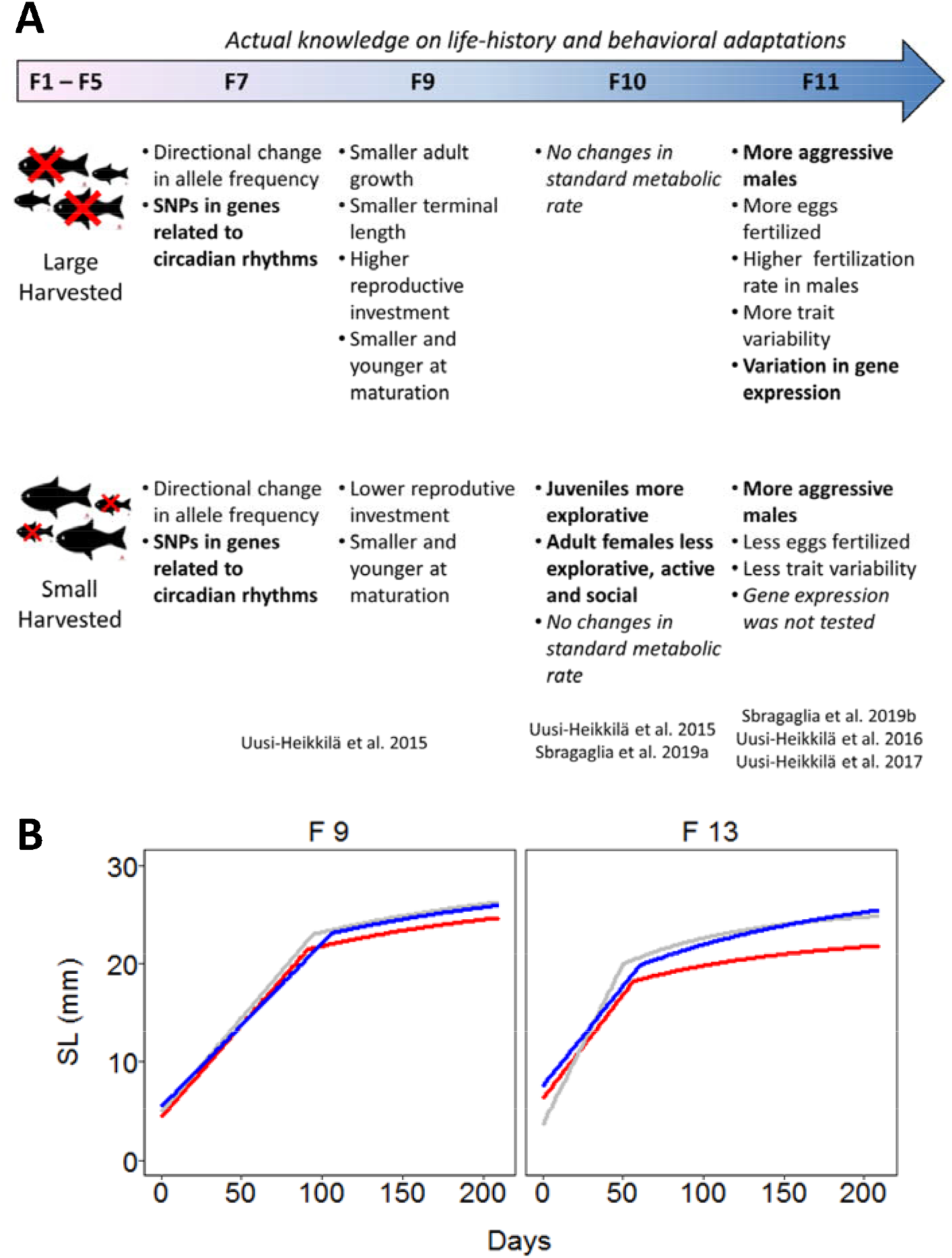
A summary of the actual knowledge on life-history and behavioral adaptations of the zebrafish selection lines used here. (A): Size-selective harvesting occurred at F_1_ – F_5_ (left). Differences in life-history and behavioral traits with respect to controls are still detectable at F_11_. The hypotheses tested in this study (F_13_) are reported on the right. The adaptations (F_7_ – F_11_) related to the hypotheses are reported in bold. (B): Fits of the Lester biphasic growth model at generations F_9_ and F_13_. Colors represent the different selection lines: large-harvested (red), control (grey) and small-harvested (blue). Adapted from Uusi-Heikkilä *et al.* (2015) and Sbragaglia *et al.* (2019b).

**Figure S2 –.**
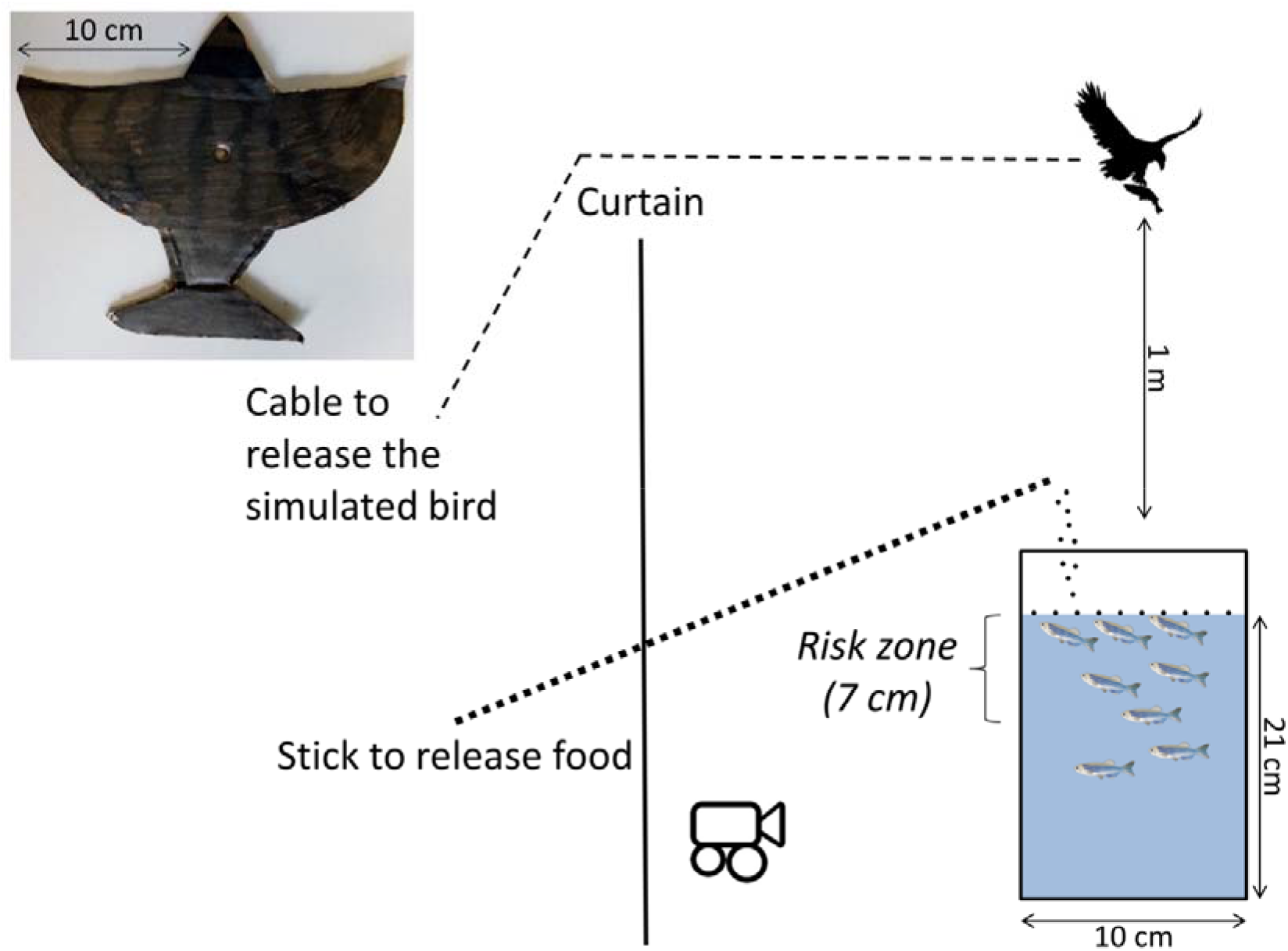
Experimental tank (10 x 30 x 25 cm) used to measure the time spent by zebrafish at the surface (risk zone = 7 cm). The tank was placed behind a curtain and a webcam was used to video record zebrafish behavior. The total experimental assay lasted 6:30 min. Recording started after the introduction of the zebrafish shoals. Food was added after 3 min and then after 30 s a simulated bird (see picture in the upper left corner) approached from above the experimental tank. The simulated bird was maintained above the tank for at about 5 seconds (See also Video S1). Zebrafish pic source: https://commons.wikimedia.org; Bird pic source: https://www.piqsels.com

**Figure S3 –.**
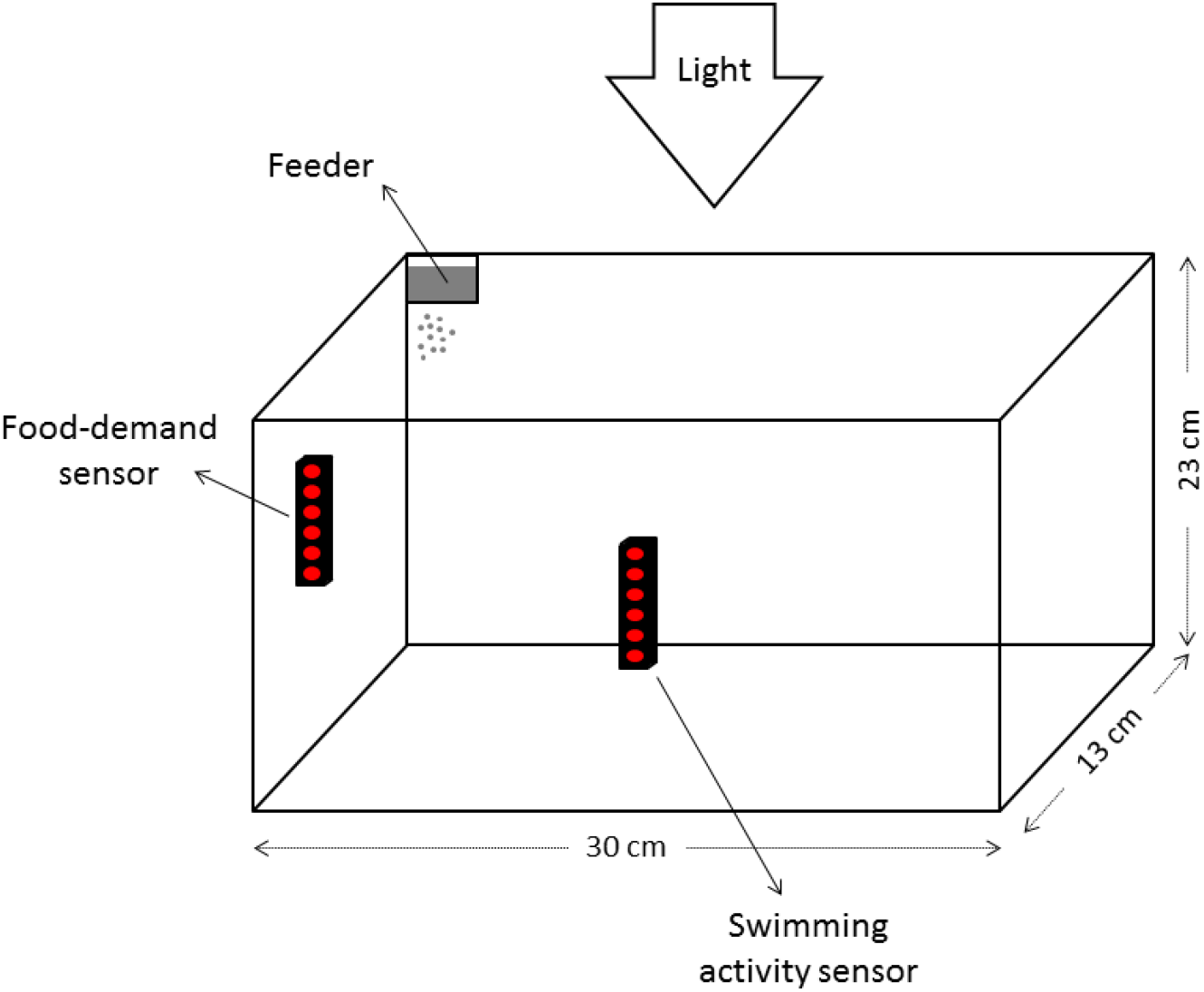
A schematic representation of the 8-L experimental aquarium (30×13×23 cm) used in this study to measure swimming and self-feeding activity rhythms in groups of 15 zebrafish. Each aquarium was equipped with aeration and mechanical and biological filters. Light was provided by white LEDs (LED Flex-strip 1043-W, Solbright, RAYTE, Murcia, Spain), with a light intensity of 435 lux at the water surface. The photoperiod was set at a 12:12 h LD cycle, and a heater (75 W Magictherm; Prodac, China) kept the water temperature at 26 ± 0.5 °C. The lights during acclimation and experimental trials were set to turn on and off at 09:00 h and 21:00 h, respectively. An infrared photocell (E3S-AD62 Omron, Kyoto, Japan) was installed in the middle of the longer side of each aquarium, 15 cm below the water surface. The sensitivity of this photocell was adjusted to detect fish at any distance from the sensor to cover the full length of the aquarium (30 cm) and measure the swimming activity. The self-feeding system consisted of another photocell located in the upper corner of the aquarium, 2.5 cm below the water surface and 1.5 cm from the side. The sensitivity of the photocell was adjusted to detect fish approaching within a range of 1 cm independently of the size range used in the experiment. The photocell was connected to a commercial feeder (Eheim, Deizisau, Germany) and when zebrafish interrupted the infrared light beam, the feeder turned on and at about 0.2% g flakes/biomass dropped into the aquarium. The feeder was placed in the same area, 10 cm from the sensor (for more details see del Pozo, Sanchez-Ferez & Sanchez-Vazquez 2011). Swimming and self-feeding activities were recorded and stored in a computer every 10 min (for more details see del Pozo, Sanchez-Ferez & Sanchez-Vazquez 2011).

**Figure S4 –.**
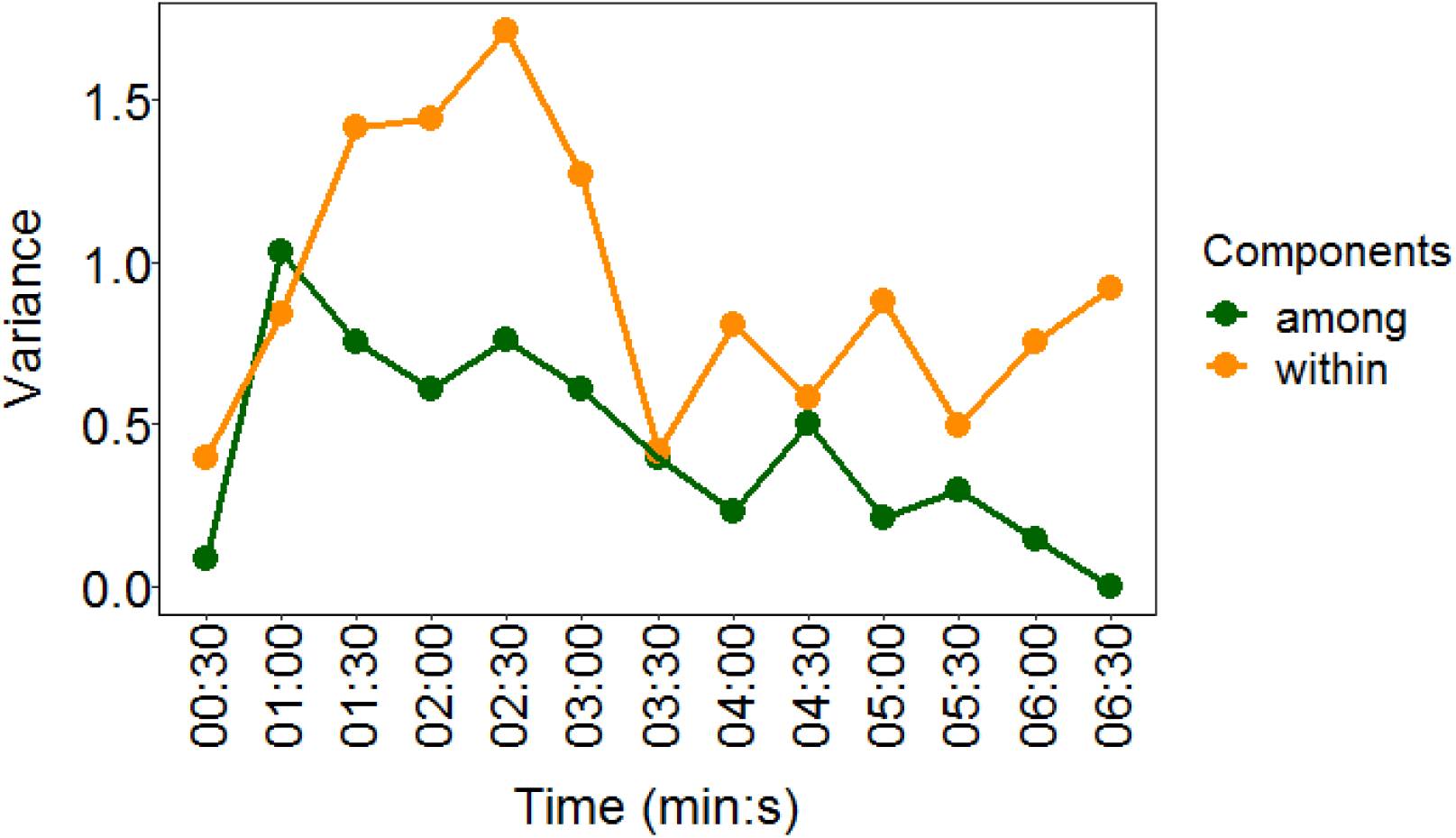
Partitioning of variance components in among group variance (see also V_g_ in Table S1) and within group variance throughout the experimental assay. Variance components are conditional on fixed effects associated to the different selection lines (*N* = 36).

**Figure S5.**
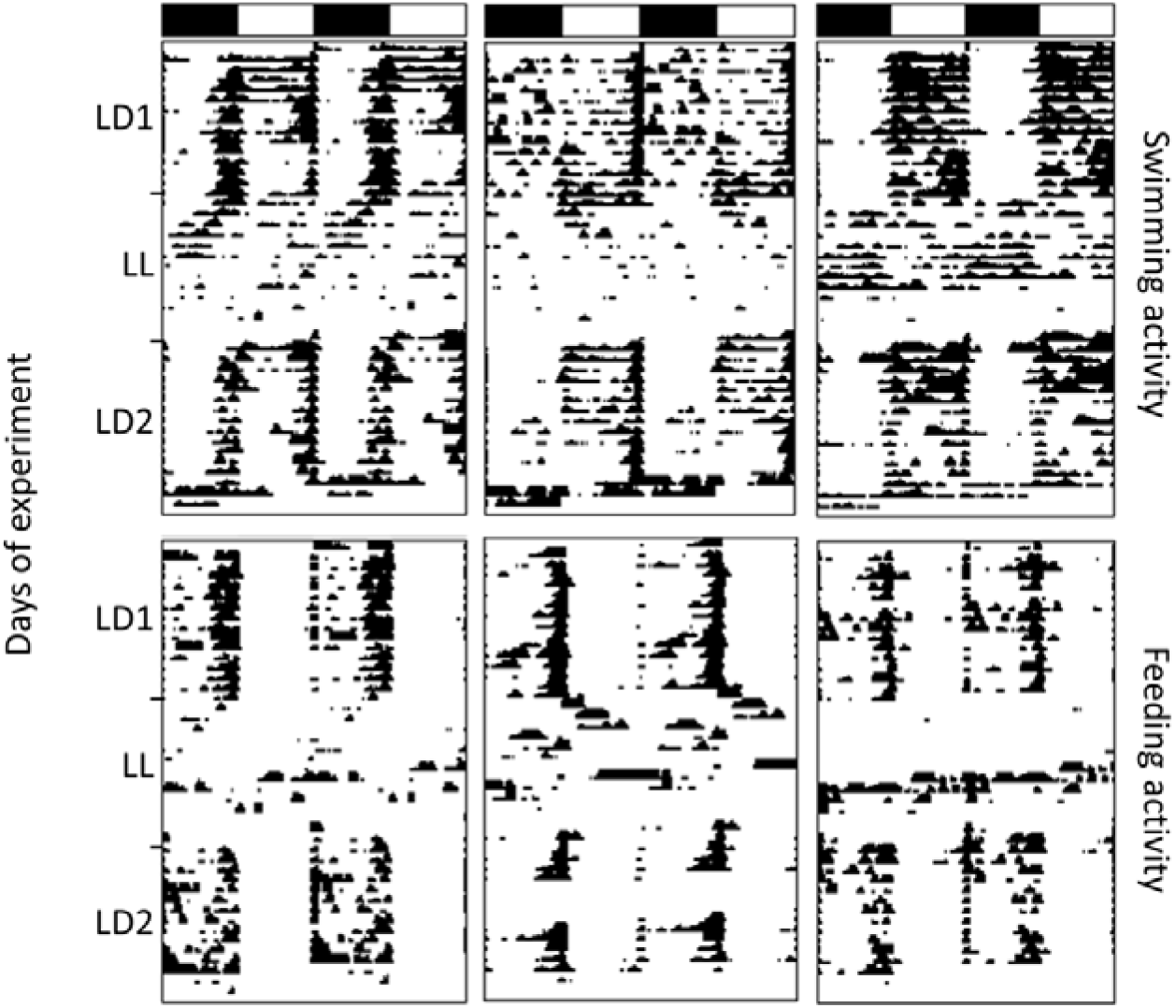
Six representative double plotted (time scale 48 h) actograms (one for each selection line) representing locomotor and self-feeding activity over 38 days of experiment (left: large-harvested line; middle: control line, right: small-harvested line). Swimming and self-feeding activity of zebrafish are indicated by the height of black areas that represent the number of infrared light beam interruptions/10 min. Open/filled bars at top represent a 12-h light/12-h darkness photoperiod for the first (LD1) and second (LD2) light-darkness trial, while during LL zebrafish shoals were exposed to constant dim light.

**Figure S6 –.**
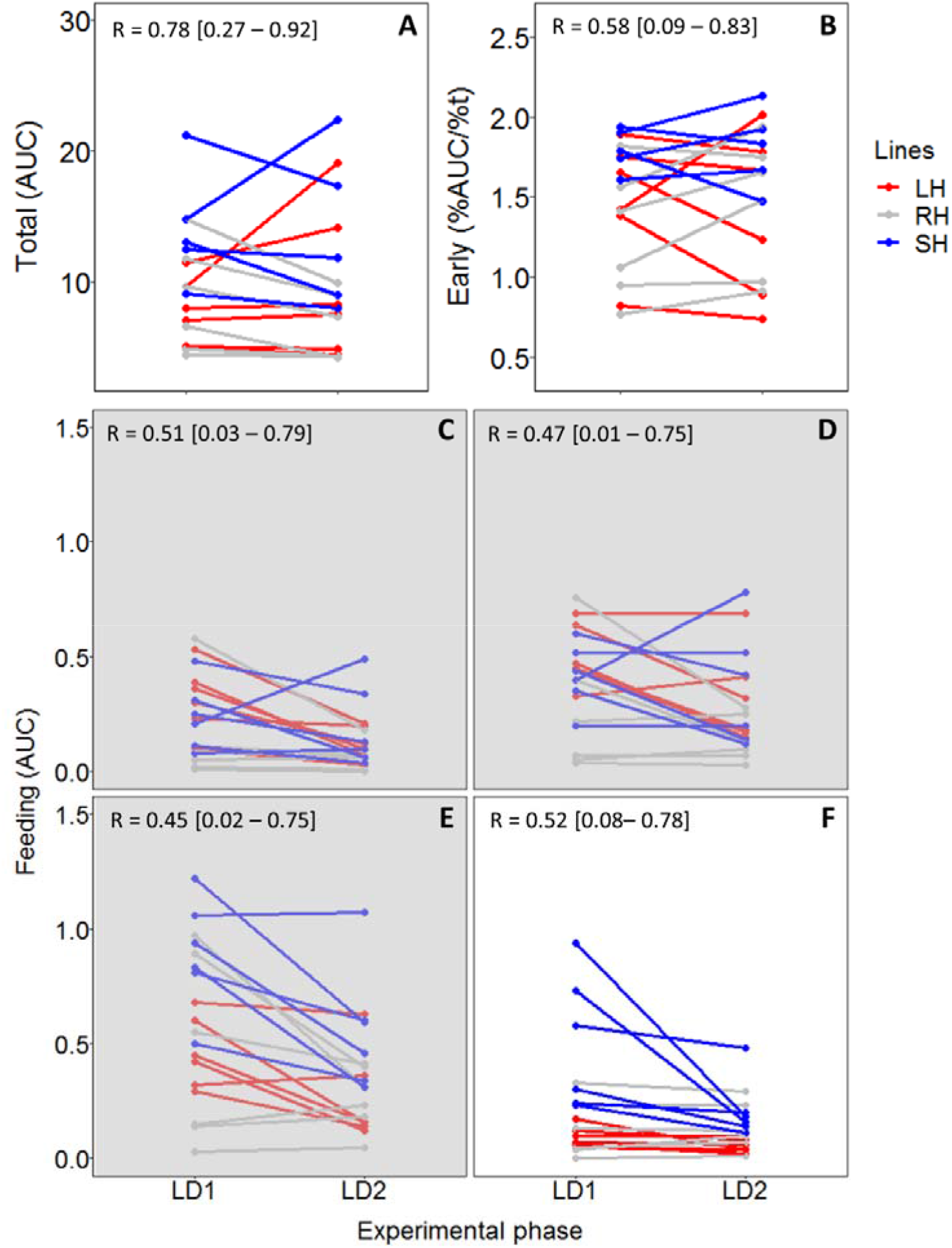
Total swimming activity during light hours (area under the waveform curve, AUC; A) and early daily activity (B; percentage of activity during the first four hours of light) are reported together with self-feeding activity (C-F; area under the waveform curve, AUC) during the last three hours of scotophase (grey shadow; C = 06:00 - 07:00; D = 07:00 - 08:00; E = 08:00 - 09:00) and the first hour of light (F = 09:00 - 10:00). Values are reported for each group at LD1 and LD2 together with the repeatability score and confidence intervals. The different colors represent the selection lines: red for large-harvested; grey for control and blue for small-harvested (N between 5 and 6; see table 1 for more details).

